# The discovery, distribution and evolution of viruses associated with *Drosophila melanogaster*

**DOI:** 10.1101/021154

**Authors:** Claire L. Webster, Fergal M. Waldron, Shaun Robertson, Daisy Crowson, Giada Ferrari, Juan F. Quintana, Jean-Michel Brouqui, Elizabeth H. Bayne, Ben Longdon, Amy H. Buck, Brian P. Lazzaro, Jewelna Akorli, Penelope R. Haddrill, Darren J. Obbard

## Abstract

*Drosophila melanogaster* is a valuable invertebrate model for viral infection and antiviral immunity, and is a focus for studies of insect-virus coevolution. Here we use a metagenomic approach to identify more than 20 previously undetected RNA viruses and a DNA virus associated with wild *D. melanogaster*. These viruses not only include distant relatives of known insect pathogens, but also novel groups of insect-infecting viruses. By sequencing virus-derived small RNAs we show that the viruses represent active infections of *Drosophila.* We find that the RNA viruses differ in the number and properties of their small RNAs, and we detect both siRNAs and a novel miRNA from the DNA virus. Analysis of small RNAs also allows us to identify putative viral sequences that lack detectable sequence similarity to known viruses. By surveying >2000 individually collected wild adult *Drosophila* we show that more than 30% of *D. melanogaster* carry a detectable virus, and more than 6% carry multiple viruses. However, despite a high prevalence of the *Wolbachia* endosymbiont—which is known to be protective against virus infections in *Drosophila*—we were unable to detect any relationship between the presence of *Wolbachia* and the presence of any virus. Using publicly available RNA-seq datasets we show that the community of viruses in *Drosophila* laboratories is very different from that seen in the wild, but that some of the newly discovered viruses are nevertheless widespread in laboratory lines and are ubiquitous in cell culture. By sequencing viruses from individual wild-collected flies we show that some viruses are shared between *D. melanogaster* and *D. simulans*. Our results provide an essential evolutionary and ecological context for host-virus interaction in *Drosophila*, and the newly reported viral sequences will help develop *D. melanogaster* further as a model for molecular and evolutionary virus research.

**Data Availability:** All of the relevant data can be found within the paper and its Supporting Information files, with the exception of raw metagenomic sequence data which are deposited at NCBI Sequence Read Archive (SRP056120), and sequence data which are deposited at Genbank (KP714070-KP714108, KP757922-KP757936 and KP757937-KP757993)

## Introduction

Viral infections are universal, and virus-mediated selection may play a unique role in evolution [1]. Viruses are also responsible for highly pathogenic diseases, and the detection, treatment, and prevention of viral disease are important research goals. The model fly, *Drosophila melanogaster*, provides a valuable tool to understand the biology of viral infection [e.g. 2,3] and antiviral immune responses in invertebrates [reviewed in 4,5], and the interaction between viruses and their vectors [e.g. 6]. *Drosophila* has also helped elucidate the role of RNA interference (RNAi) as an antiviral defence [e.g. 7,8], and has shown that endosymbiotic *Wolbachia* can protect against viruses [9,10]. Recently, the Drosophilidae have been used to address important questions in virus evolution, including determinants of host-range and disease emergence [11-13]. However, although *Drosophila* virus research has a long history, few *D. melanogaster* viruses are known in the wild [4,14], and experiments using non-natural *Drosophila* pathogens may bias our understanding of immune function and its evolution [e.g. 12].

Following the discovery of Sigma Virus in *D. melanogaster* (DMelSV, Rhabdoviridae; reviewed in [15]), classical virology surveys in the 1960s and 1970s uncovered Drosophila C Virus (DCV, Dicistroviridae), Drosophila A Virus (DAV, related to Permutotetraviridae), Drosophila X Virus (DXV, Birnaviridae), DFV (Reoviridae), DPV and DGV (unclassified) in *D. melanogaster* [reviewed in 4,14]. Subsequent transcriptomic studies of *D. melanogaster* identified *D. melanogaster* Nora Virus (unclassified Picornavirales; [16]), and analyses of small RNAs from *D. melanogaster* cell culture [17] identified Drosophila Totivirus, American Nodavirus (closely related to Flockhouse Virus), and *Drosophila* Birnavirus (closely related to DXV). However, only four of these viruses have been isolated from wild flies, have genome sequences available, and are available for experimental study. These include DCV [18], DMelSV [19], DAV [20], and Nora Virus [16], while DXV is reported to be a cell culture contaminant [14,21]. Of these four, only DMelSV has been widely studied in the field [15,22,23].

Our limited knowledge of *D. melanogaster*’s natural viruses reflects a historically tight research focus on viruses with direct medical and economic impact. While high-throughput ‘metagenomic’ sequencing has broadened our knowledge of viral diversity in general [reviewed in 24,25], most studies focus on vertebrate faeces [26,27], potential reservoirs of human and livestock disease [28-30], or crop plants [e.g. 31]. Relatively few studies have performed metagenomic virus discovery in invertebrates [for a review, see 32]. Recently, a large survey identified an exceptional and unsuspected diversity of negative sense RNA viruses associated with arthropods, suggesting that we may only have been scratching the surface of viral diversity [33]. However, aside from some lepidopteran pests [34] and hymenopteran pollinators [35], we still know little about the biology of most invertebrate viruses.

Although we have much to learn about *Drosophila* viruses, experiments using both natural and non-natural *Drosophila* pathogens have given us a better understanding of viral infection and immunity in *Drosophila* than in any other invertebrate [2,5]. DCV, DXV and Nora Virus have all been used to study the molecular biology of host-virus interaction [e.g. 7,36-39], and classical genetic approaches have elucidated the basis of host resistance to DCV and DMelSV [40-45]. Many insect viruses—notably Cricket Paralysis Virus, Flock House Virus (from beetles), Sindbis Virus (mosquito-vectored), Vesicular Stomatitis Virus (mosquito-vectored), and Invertebrate Iridovirus 6 (from mosquitoes)—have helped characterise the roles of the RNAi, IMD, Toll, autophagy, and Jak-Stat pathways in antiviral immunity [reviewed in 5]. These studies show that *Drosophila* has a sophisticated and effective antiviral immune response, and both molecular [e.g. 12] and population genetic [46-48] studies suggest that this immune system may be locked into an evolutionary arms race with viruses. However, until we understand the diversity, distribution or prevalence of viral infection in *D. melanogaster*, it is hard to put these results into their evolutionary or ecological context.

Here we use a metagenomic approach to identify more than 20 novel viruses associated with *D. melanogaster*, including the first DNA virus. Based on the presence of virus-derived 21nt small RNAs (which are characteristic of an antiviral RNAi response in *Drosophila* [49]), we argue that these sequences represent active viral infections. Using a survey of individual wild-collected flies we give the first quantitative estimates of prevalence for 15 different viruses in *D. melanogaster* and its close relative *D. simulans*, and rates of co-occurrence with the *Wolbachia* bacterial endosymbiont. In addition, by examining publicly available RNA datasets we catalogue the presence of these viruses in *D. melanogaster* stock lines and cell culture. Our results provide an unprecedented insight into the virus community of *Drosophila*, and thereby provide the evolutionary and ecological context needed to develop *Drosophila* as a model for virus research.

## Results and Discussion

### There are many virus-like sequences associated with wild *Drosophila*

We used metagenomic sequencing of ribosome-depleted total RNA to identify virus-like sequences in five large collections of wild-caught adult Drosophilidae. Three collections (denoted E, K, and I) were sampled from fruit baits in Kilifi (Kenya) and Ithaca (New York, USA). The aliquots pooled for total RNA sequencing represented around 2000 individuals of *D. melanogaster, D. ananassae*, *D. malerkotliana*, and *Scaptodrosophila latifasciaeformis*. Two collections (S and T) were sampled from fruit baits in southern England, and aliquots pooled for total RNA sequencing represented around 3000 *D. melanogaster* (<1% other *Drosophila*). In total 0.5% of all reads mapped to DAV, DCV, DMelSV, and Nora Virus (supporting information S1 and S2). Viral read numbers varied dramatically between samples, with DCV absent from pool EIK and DAV absent from pool ST (verified by qPCR; supporting information S3). Only 3 reads mapped to DXV, and no reads mapped to Drosophila Totivirus, Drosophila Birnavirus, or American Nodavirus (S1). As DXV is reported to be a cell culture contaminant [14], and the other unmapped viruses were described from cell culture [17], this suggests that these viruses may be rare in wild flies.

To discover novel viruses, we assembled all RNA-seq reads *de novo* using Trinity [50] and Oases [51], and used BLAST [52] against the Genbank protein reference sequences [53] to identify virus-like sequences. Raw *de novo* metagenomic contigs are provided in supporting information S5. Virus-like contigs were supplemented with PCR and targeted Sanger sequencing to improve completeness, and in total we identified more than 20 partial viral genomes. Those sequences which could be unambiguously associated with *D. melanogaster* rather than other *Drosophila* species were provisionally named according to collection locations. Based on sequence similarity, these ‘BLAST-candidate’ viruses included two Reoviruses (‘Bloomfield Virus’ and ‘Torrey Pines Virus’), three Flaviviruses (‘Charvil Virus’, and two others), a Permutotetravirus (‘Newfield Virus’), a Nodavirus (‘Craigie’s Hill Virus’), a Negevirus, a Bunyavirus, two Iflaviruses (‘La Jolla Virus’ and ‘Twyford Virus’), two Picorna-like viruses (‘Thika Virus’ and ‘Kilifi Virus’), a virus related to Chronic Bee Paralysis Virus (‘Dansoman Virus’), a virus related to Sobemoviruses and Poleroviruses (‘Motts Mill Virus’), six Partitiviruses, and a Nudivirus (‘Kallithea Virus’). These novel viruses constituted a further 3.3% of RNA-seq reads, taking the viral total to 3.8% of all reads (S1). Further details of the new viruses are given in Table 1 and supporting information S6, and virus sequences have been submitted to Genbank as KP714070-KP714108 and KP757922-KP757936.

**Table 1:**
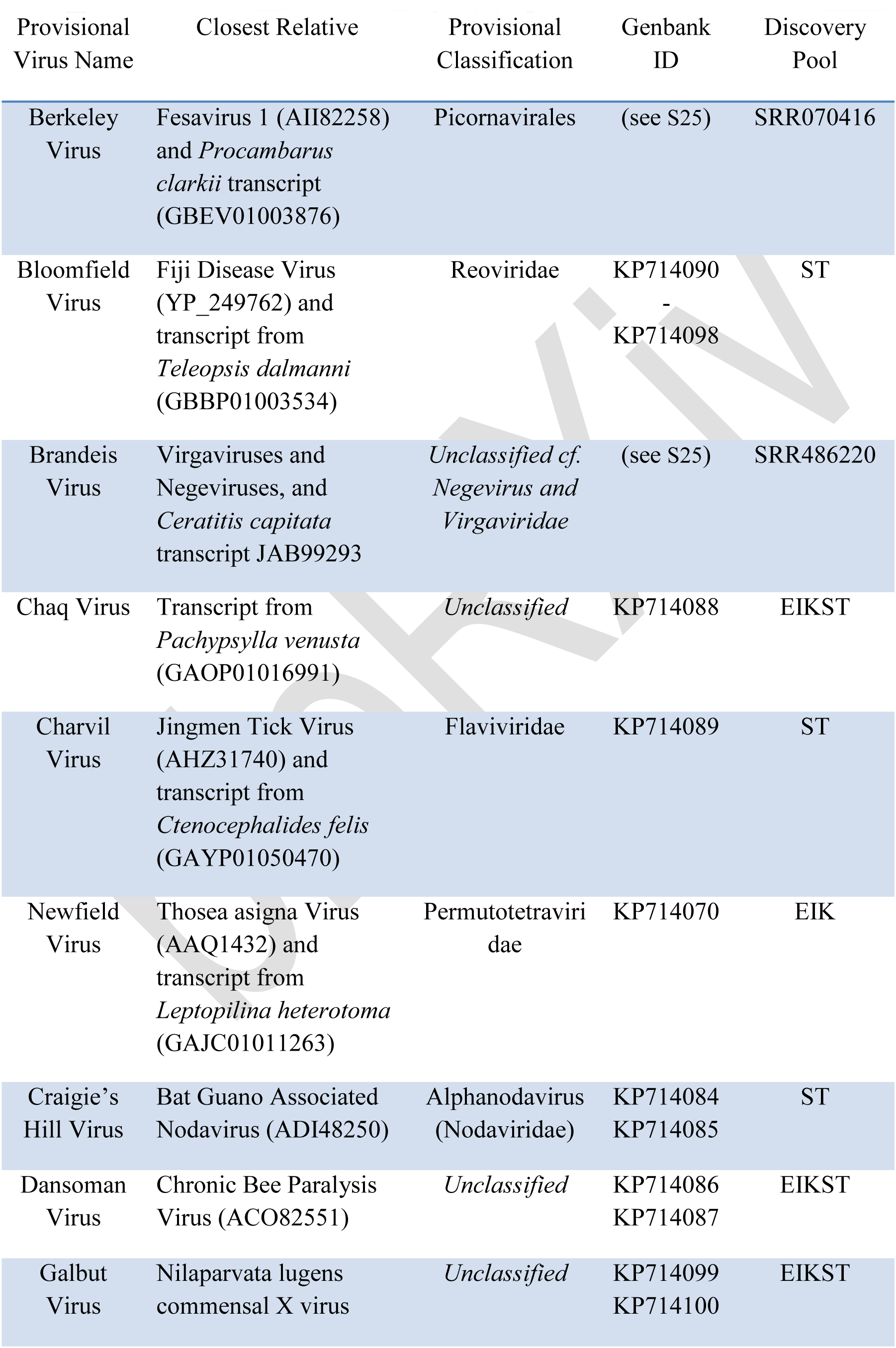
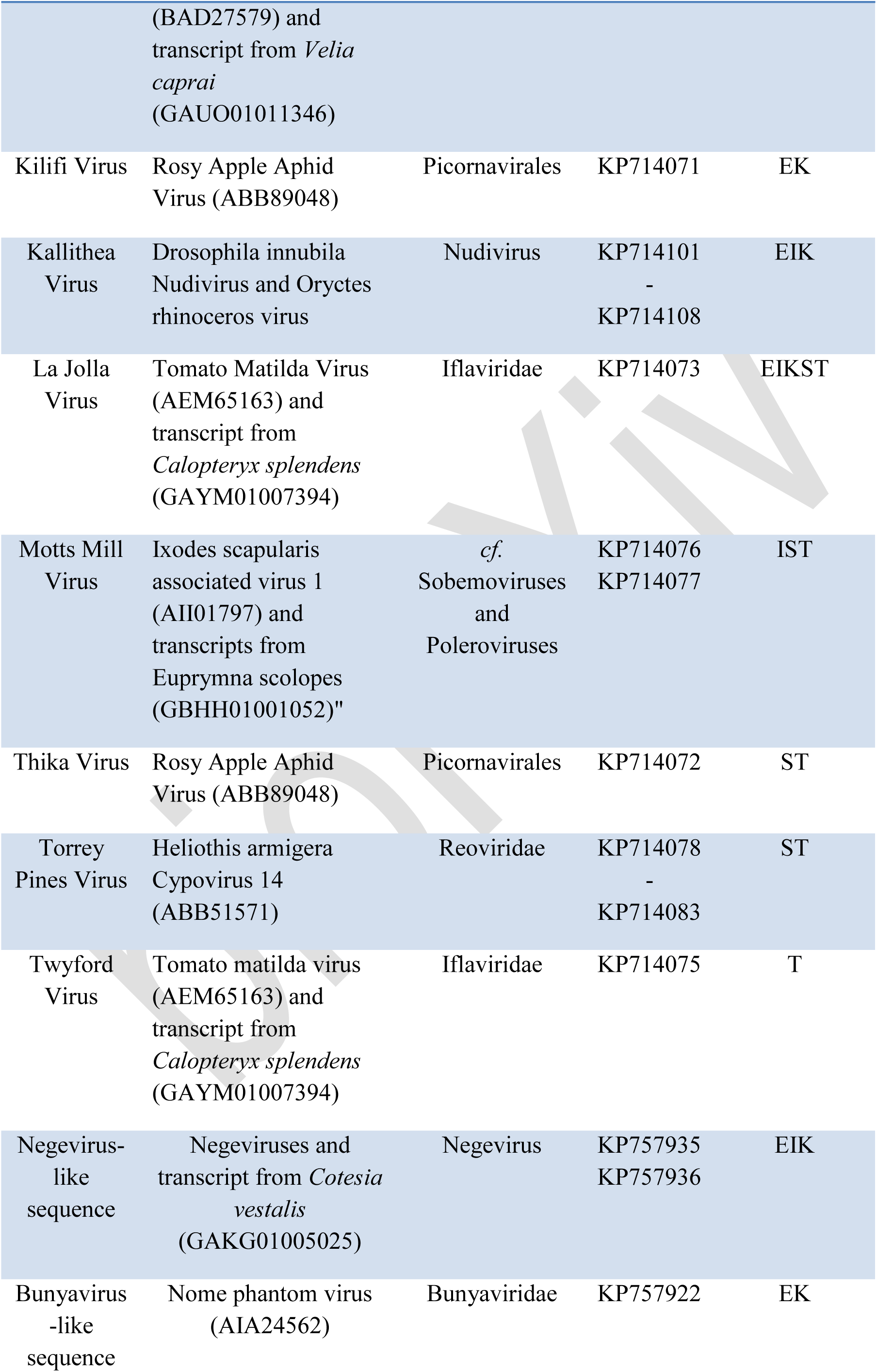
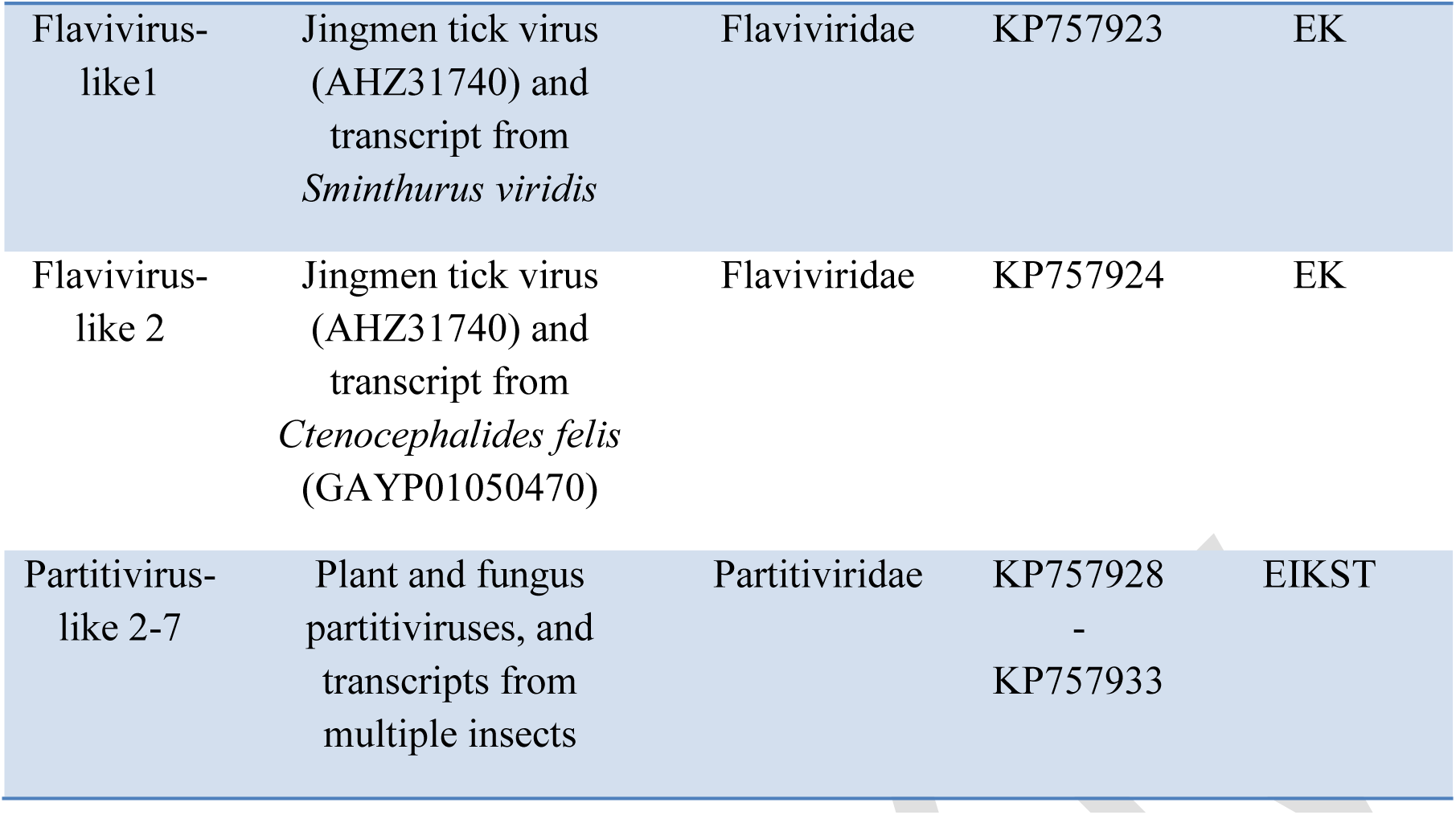
New *D. melanogaster*-associated Viruses

To place the virus sequences in a phylogenetic context, we subjected conserved regions to phylogenetic analysis along with known viruses and un-curated viral sequences from the NCBI Transcriptome Shotgun Archive [e.g. 54]. Kallithea Virus, a DNA virus, was closely related to Nudiviruses from *Drosophila innubila* [55] and the beetle *Oryctes rhinoceros* [56]. Most of the new RNA viruses were relatives of known or suspected insect pathogens (Table 1, supporting information S7). For example, based on polymerase sequences, Torrey Pines Virus is distantly related to *Aedes pseudoscutellaris* reovirus and several lepidopteran Cypoviruses, while Dansoman Virus is related to Chronic Bee Paralysis Virus. Others were distantly related to arthropod viruses, but were close to un-curated transcriptome sequences (Figure 1). For example, Bloomfield Virus is closely related to transcriptome sequences from the flies *Delia antiqua* and *Teleopsis dalmanni* (63% AA identity in the replicase), and distantly related to *Nilaparvata lugens* reovirus. Similarly, Motts Mill Virus is related to the recently described *Ixodes scapularis* (tick) associated viruses 1 and 2 [57], but is closer to un-curated transcriptomic sequences from three bees and the bobtail squid *Euprymna scolopes*. Together these sequences appear to represent a novel clade related to plant Sobemoviruses and Poleroviruses (Figure 1). Strikingly, the six unnamed Partitivirus-like polymerases were not related to any known insect pathogens (known Partitiviruses are pathogens of plants and fungi), but were related to un-curated sequences from arthropod transcriptomes—again suggesting a novel lineage of insect viruses (Figure 1).

**Figure 1:**
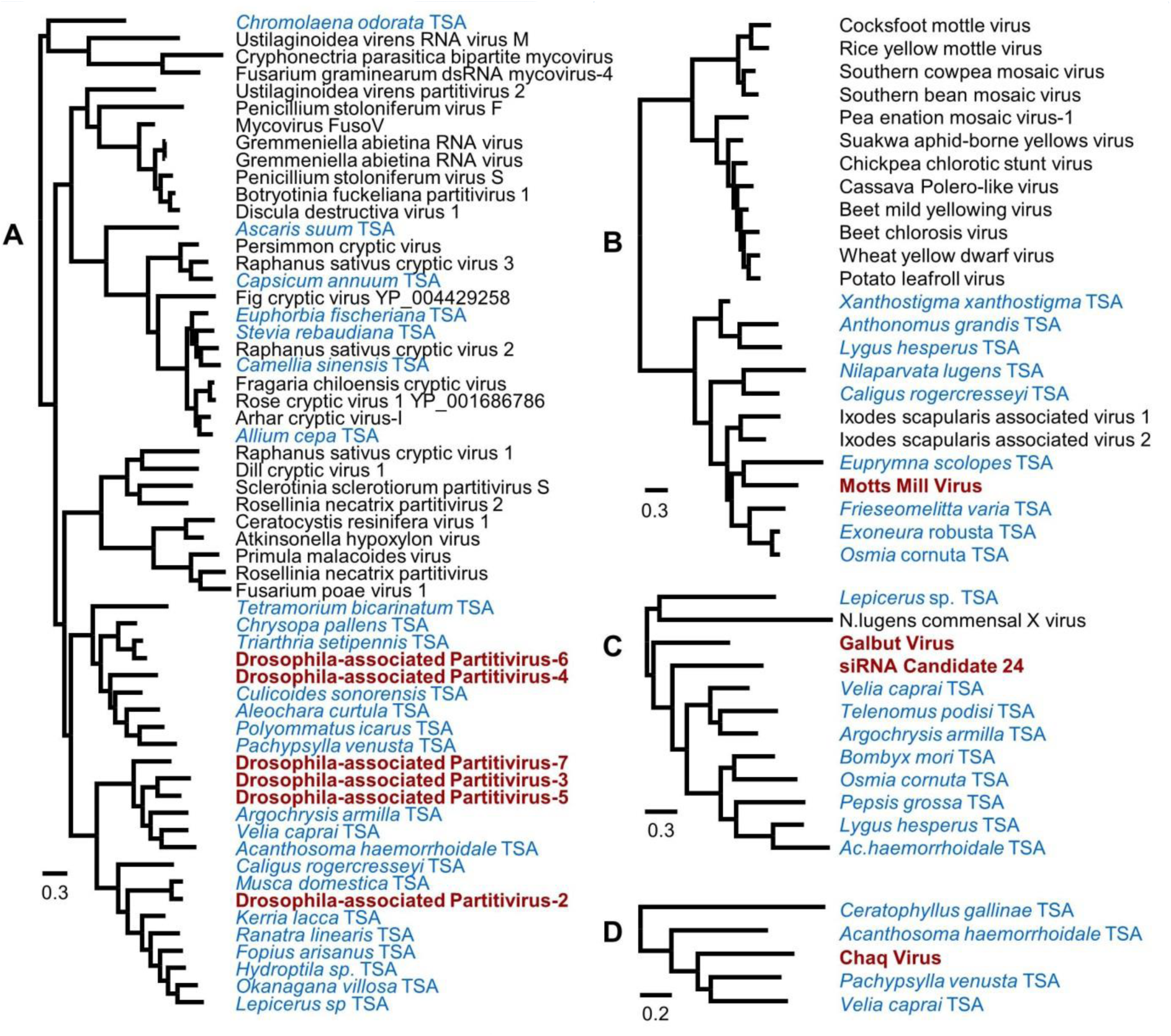
Phylogenetic analysis of viruses not closely related to known insect pathogens. Although the majority of viruses were closely related to known insect pathogens (Table 1, supporting information S7), a minority were not, and may represent novel insect-infecting lineages. These include (tree A) six sequences similar to the Partitiviridae, (tree B) the BLAST-candidate Motts Mill Virus, distantly related to Luteoviridae and Sobemoviruses, (tree C) the siRNA-candidate Galbut Virus, and (tree D) the siRNA-candidate Chaq Virus. Trees are mid-point rooted maximum clade credibility trees inferred from the Bayesian posterior sample, and the scale is given in amino-acid substitutions per site. Node support and sequence accession identifiers are given in supporting information S7. *Drosophila*-associated sequences discovered here are shown in red, Transcriptome Shotgun Archive sequences in blue. The Partitivirus-like sequences are all closely related to sequences present in insect transcriptome datasets, suggesting they may form a novel clade of insect pathogens. Motts Mill Virus similarly clusters with invertebrate transcriptome sequences and two recently reported viruses from *Ixodes* ticks [57], again suggesting a new clade of pathogens. Neither Chaq Virus nor Galbut Virus has high sequence similarity to known viruses, but both also cluster with invertebrate transcriptome-derived sequences. These may represent new virus lineages, or be weakly conserved genes in a known virus group. Alignments with MrBayes command blocks are provided in supporting information S8, and maximum clade credibility trees are provided in supporting information S9.

The close relationship between the newly identified virus-like sequences and known arthropod viruses suggests that they are likely to be *Drosophila* pathogens, and some may be previously described viruses that lack sequence data. For example, Kilifi Virus or Thika Virus (related to the Picornavirales; S7) may correspond to DPV—which was reported to have a 25-30nm particle and infect gut tissues [58]. Similarly, either Torrey Pines Virus or Bloomfield Virus could correspond to Drosophila F Virus [14], Drosophila K Virus or other reoviruses [see 21]. However, as those viruses were described only from capsid morphology, density, and serology, it would be challenging to conclusively link them with the novel sequences presented here.

### Genomic data and small RNAs imply active viral infection

Although phylogenetic analyses show that these sequences are viral in origin, they may represent Endogenous Viral Elements integrated into the *Drosophila* genome (‘fossil’ viruses, or EVEs [59]), or they may derive from gut contents or surface contamination rather than active infections. To exclude the possibility that these sequences represent EVEs segregating in *D. melanogaster*, we mapped the raw genomic reads from 527 distinct *D. melanogaster* genomes [60] to our set of BLAST-candidate viruses, and confirmed that no genome mapped at a rate high enough to be consistent with a genomic copy of any virus in that individual. As this test for EVE status was not possible for other *Drosophila* species, only sequences associated with *D. melanogaster* were named as viruses, and others (which could potentially represent EVEs in other taxa) were denoted ‘virus-like’.

To test if these sequences derive from active viral infections, we additionally sequenced all 17-29 nt small-RNAs from the EIK and ST pools, reasoning that virus-like sequences will only be processed into 21 nt siRNAs (viRNAs, derived from replicative intermediates by Dicer-2) if they represent active infections within host cells [49,61]. In total *ca.* 7% of all 17-29nt small-RNAs derived from DAV, DCV, DMelSV, and Nora Virus, and *ca.* 9% derived from the new ‘BLAST-candidate’ viruses. As expected, for most viruses the viRNA size distribution was tightly centred on 21nt reads and included reads from both the genomic and complementary strands, consistent with active viral infections processed by antiviral RNAi in *Drosophila* (Figure 2; S11).

**Figure 2:**
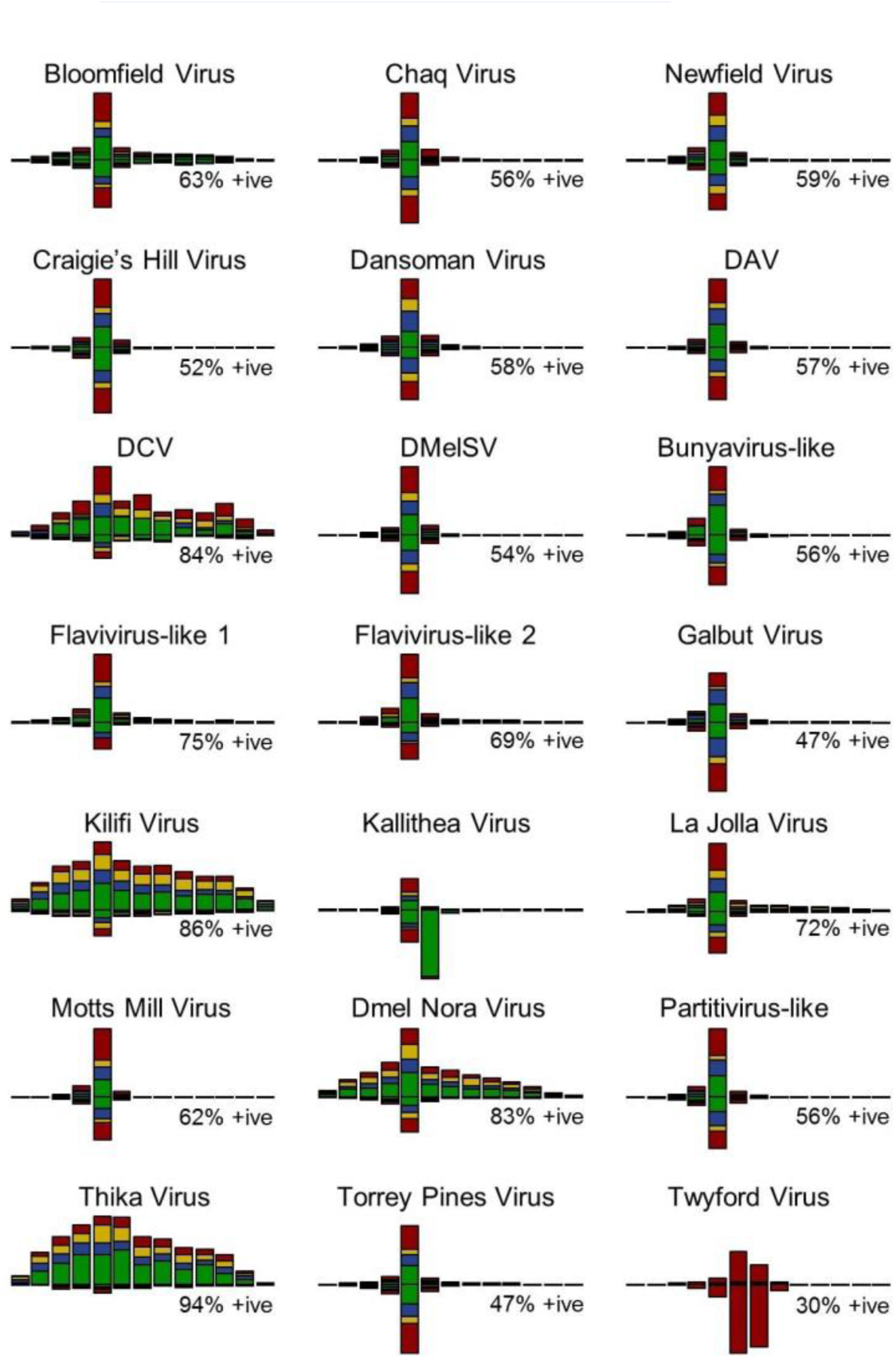
Small RNA size distribution and base composition. The size distribution and base composition of small (17-29nt) RNAs varied substantially amongst viruses. The barplots show the size distribution for those viruses with >100 small RNA reads (summed across all libraries, for a breakdown see S11). Bars plotted above the x-axis represent reads mapping to the positive strand, and bars below represent reads mapping to the negative strand. All peak at 21nt, except Kallithea Virus (a large DNA Nudivirus encoding a 22nt miRNA) and Twyford Virus (Iflavirus), which peak at 22nt. Note that DCV, Kilifi, Thika and Nora Virus show a wide shoulder up to 28nt for positive-strand reads, which may indicate a difference in small RNA processing or retention. Bars are coloured according to the proportion of reads with each 5'-base (A-green, C-blue, G-yellow, U-red), and all except Twyford Virus show the expected pattern, including a slight bias against G at the 5’ position. Counts used to plot this figure are provided in supporting information S10.

Even if viral sequences do not represent EVEs or inactive contaminants, they could instead be active infections of *Drosophila*-associated microbiota rather than *Drosophila*. However, the 21nt viRNAs observed for the majority of these viruses are inconsistent with viral infection of likely parasites or parasitoids such as hymenoptera, chelicerata or nematodes, which have predominantly 22nt viRNAs [e.g. 62,63]. And, while 21 nt viRNAs could derive from viral infections of some fungi [e.g. 64] or from eukaryotes with uncharacterised antiviral RNAi, in most cases the phylogenetic position of these viruses, high read numbers, and/or their appearance in laboratory fly stocks and cell culture (below) argue in favour of *Drosophila* as the host.

### Small RNAs are markers for novel virus-like sequences

In addition to the viral candidates identified by BLAST, we reasoned that contigs which lack BLAST similarity to reference sequences, but which display a signature of Dcr-2 processing (high levels of 21-23 nt siRNAs and low levels of 25-29 nt piRNAs), may also be viral in origin (Figure 3; [see e.g. 17,64]). Using these small RNA criteria we identified a list of ‘siRNA-candidate’ contigs that are potentially viral in origin, but could not be placed within a phylogeny of known viruses. Several siRNA-candidate viruses identified in this way were subsequently attributable as fragments of the BLAST-candidate viruses by other means (below). Of those that could not be attributed to viral genomes, two were provisionally named (Chaq Virus and Galbut Virus) and the remaining 57 contigs were submitted to Genbank as uncultured environmental virus sequences (KP757937-KP757993). These unnamed siRNA-candidate viruses contribute 0.2% of all RNAseq reads, and 1% of 17-29nt small RNAs in the EIK and ST pools (S1). As with the BLAST-candidate viruses, these siRNA-candidate viruses were absent from *D. melanogaster* genomic reads and their siRNAs were predominantly 21nt in length and derived from both strands (S16), again suggesting that they represent active viral infections of *Drosophila*. Although the siRNA-candidate viruses displayed no strong BLAST similarity to known viruses, Galbut Virus and siRNA-candidate 24 display weak similarity to *Nilaparvata lugens* Commensal X Virus (a satellite virus with unknown helper [65]), and Galbut and Chaq viruses appear to be related to un-curated sequences present in a diverse set of arthropod transcriptome shotgun datasets (Figure 1 and Table 1; S6 and S7).

**Figure 3:**
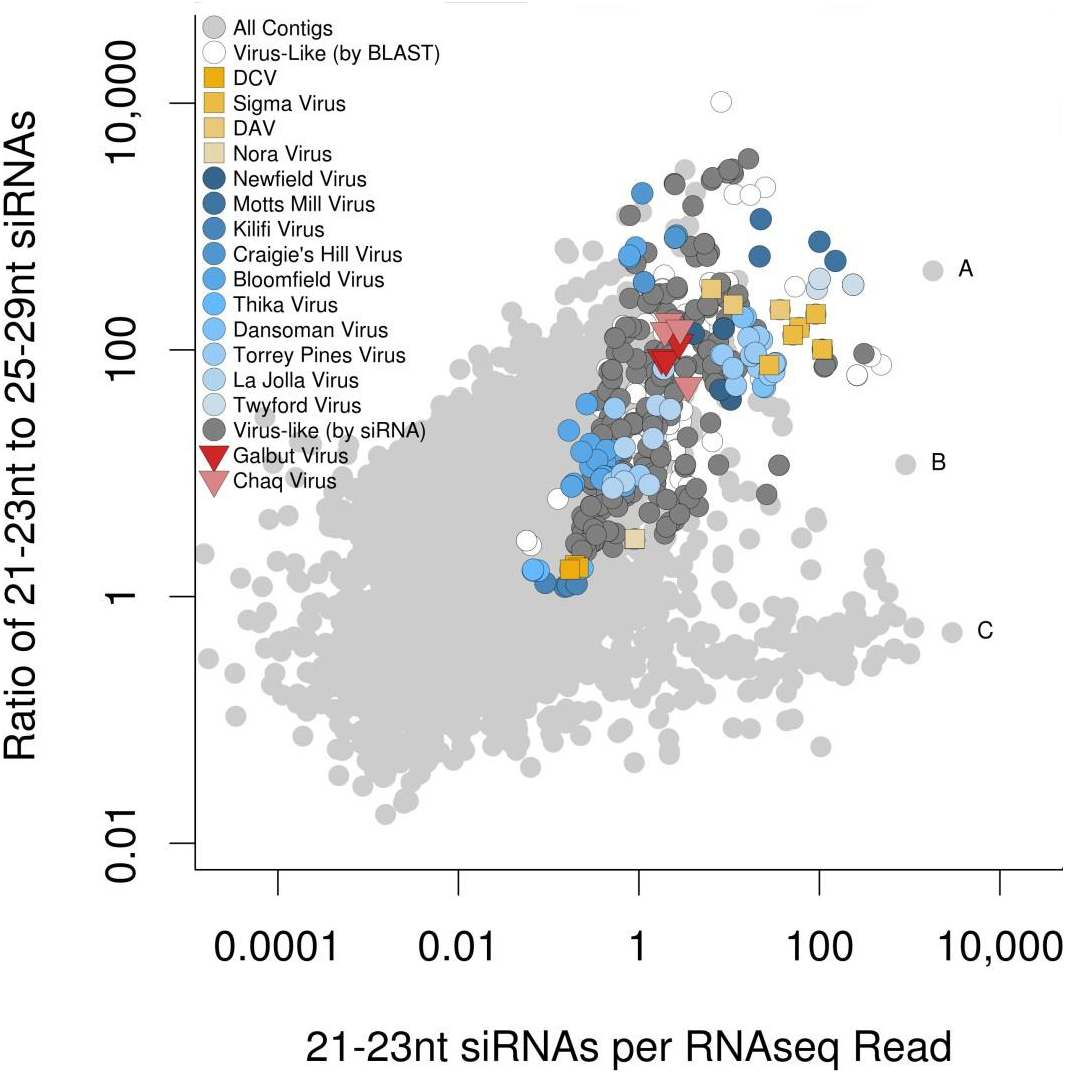
Identification of virus-like sequences via small RNAs. Small RNA and RNA-seq reads were mapped to all *de novo* Trinity and Oasis contigs (regardless of origin), and the numbers of 21-23nt RNA reads (viRNA-like) and 25-29nt RNA reads (piRNA-like) were recorded relative to the number of RNAseq reads. For small RNAs all four of the mixed ST and EIK pools were combined, while for RNAseq data a single rRNA-depleted non-normalised mixed pool was used. Contigs with at least one small RNA read are shown in pale grey, and contigs are classifiable as more ‘virus-like’ toward the top right as the total number of siRNAs increases and the ratio of siRNA:piRNA reads increases. As expected, DAV, DCV, Nora Virus, and DMelSV (yellow squares) have large numbers of siRNAs and few piRNAs, as do named BLAST-candidate viruses (blue circles), and other contigs with sequence similarity to known viruses (white circles). Based on the sector delimited by these viruses, we identified contigs with no BLAST hits in either Genbank refseq_rna or refseq_protein that we designate ‘siRNA candidate viruses’ (dark grey). Two of these were given provisional names (Galbut Virus and Chaq Virus: red triangles). Note that each virus may appear more than once in the figure, if more than one contig derives from that virus. Outliers include (A) a miRNA-bearing fragment from Kallithea virus, (B) unknown sequence with no strong blast hit, and (C) a Jockey-like retrotransposon, expected to be a source of piRNAs. The read-mapping count data are provided in supporting information S12, and raw *de novo* contigs are provided in supporting information S5.

### RNA viruses differ in the numbers and properties of small RNAs

To test whether viruses vary in their small RNA profile, we analysed small RNAs from the EIK and ST pools, and from two libraries for each of the five collections (E, I, K, S, and T), with and without ‘High Definition’ ligation adaptors designed to reduce ligation bias [e.g. 66]. Overall, the number of viRNA reads per RNAseq read varied substantially among viruses (‘viRNA ratio’; Figure 3 and S18), with Twyford Virus and Motts Mill Virus giving rise to more than a 1000-fold more 21-23nt viRNAs than DCV, and DMelSV giving rise to nearly 7000-fold more (S18). For viruses present in both EIK and ST, the viRNA ratios were highly correlated between pools (rank correlation ρ>0.99; S18), suggesting that they are repeatable. This may reflect differences in the proportion of non-replicating viral genomes (e.g. encapsidated viruses in the gut lumen) which can contribute to RNAseq but are not actively processed by Dcr-2. Alternatively, differences could result from the action of viral suppressors of RNAi (VSRs), such as those encoded by DCV and Nora Virus [7,38].

The sizes and strand bias of small RNAs also varied substantially among viruses, although small RNA reads from the majority of RNA viruses were biased toward the positive strand (50-70%) and to 21 nt in length (Figure 2), as expected for virus-derived small RNAs in *Drosophila* [e.g. 49]. Exceptions included Nora Virus, DCV, Kilifi Virus and Thika Virus, which showed a stronger positive-strand bias (85-95% of reads) and a broad size range of positive-sense viRNAs peaking at 21nt, with a wide ‘shoulder’ from 23 to 27nt (Figure 2). This size distribution is not seen in most DCV infections of cell culture [49], and although 26-30nt viral piRNAs have been reported in OSS cell culture [17], the 25-28nt reads identified here did not display the 5' U bias expected of piRNAs (Figure 2; [67]). As these four picorna-like viruses also displayed low numbers of viRNAs (Figure 3; S2, S10, S11), this is consistent with a difference in the way that the viral genome and/or viRNAs are processed—perhaps reflecting a higher fraction of non-specific degradation products for these viruses. However, because we found that viRNA properties were reproducible across sequencing libraries (S11), and because viRNAs were sequenced from large pooled samples composed of mixed infections, these profiles cannot result from idiosyncrasies of RNA extraction or library preparation.

In contrast to the other RNA viruses, Twyford Virus displayed unusual viRNAs. Although Twyford Virus is an Iflavirus (S7) and thus has a positive sense ssRNA genome, the strand bias was strongly negative and the viRNAs peaked sharply at 22-23 nt (*cf.* 21 nt for other RNA viruses). In addition, although other viruses displayed no strong 5' base-composition bias except for a weak bias against 5' G, most Twyford Virus viRNAs were 5' U, as is seen for piRNAs (but lacking the A at position 10 expected of *Drosophila* piRNAs; S14). This bias is not due to small sample size or low sequence diversity as we saw >9 thousand unique sequences, and 3500 of those were seen more than once. Comparison with the virus genome showed the 5' U bias was driven by differential production or retention of viRNAs, rather than subsequent editing or addition. Interestingly, the 3' position was also slightly enriched for U (S13), and a substantial fraction of 3' U were non-templated, indicative of 3' uridylation as seen in *D. melanogaster* and other species in the absence of the Hen-1 methytransferase [68].

These observations suggest that Twyford Virus may not be processed by the *Drosophila* Ago2-Dcr2 pathway, and could instead represent viral infection of an unknown eukaryotic commensal. Nevertheless, potential arthropod parasites such as chelicerata have not been reported to display this pattern of viRNAs, and although the 5' U is reminiscent of the 21U piRNA of *C. elegans* and related nematodes [69], neither Rhabditid nor Tylenchid nematodes could be detected by PCR in individual wild-caught *D. melanogaster* carrying Twyford Virus. Similarly, while 22nt 5'-U small RNAs are known from the filamentous fungus *Neurospora* [70], those are derived from the host genome and are not associated with viral infection. Thus, although speculative, if these 22-23U viRNAs cannot be explained by a non drosophilid host, then it is possible that they reflect a previously unrecognised tissue-specific phenomenon in *Drosophila*, or one associated with the action of a novel suppressor of RNAi (e.g. suppression of Hen-1). The siRNA-Candidate 14 (KP757950) shows a similar pattern (22-23nt peak in viRNAs,5' U), suggesting is forms part of the same virus and/or is processed in the same way (S16).

### Kallithea Virus is targeted by RNAi and encodes a miRNA

We also identified many 21 nt viRNAs widely-dispersed around the Kallithea Virus genome. This is consistent with an antiviral RNAi response against this DNA Nudivirus, as has been previously reported for Invertebrate Iridovirus 6 artificially infecting *D. melanogaster* [71]. As DNA viruses often encode miRNAs [72,73], and miRNAs have been implicated in the establishment of latency in Heliothis zea Nudivirus [74], we screened small RNAs from Kallithea Virus for potential virus-encoded miRNAs. One 22nt RNA sequence was highly abundant (S14), and is predicted to represent the 5' miRNA from a pre-miRNA-like hairpin (miRDeep2 [75]; S20). The predicted mature 5' miRNA (AUAGUUGUAGUGGCAUUAAUUG) represented >35% of all small RNAs derived from Kallithea Virus, while the 3' RNA ‘star’ sequence represented 0.3% of reads. This sequence was less highly represented, relative to 21nt viRNAs from Kallithea Virus, in an oxidised library, consistent with the absence of 2'O-methylation at the 3' end, as expected for miRNAs in *Drosophila* ([76]; S11). The seed region displays no obvious similarity to known miRNAs (although positions 5-17 are similar to dme-miR-33-5p), but a survey of potential binding sites in *D. melanogaster* using miRanda 3.3a [77] identified 522 genes with at least one potential binding site in the 3'-UTR (miRanda score ≥150). These were highly enriched for Gene Ontology terms that might be associated with viral function (including, amongst others: Regulation of Gene Expression, Cell development, mRNA binding, Plasma Membrane; S22). This miRNA could alternatively regulate virus gene expression [e.g. 74], and 21 potential binding sites were identified in the Kallithea Virus genome. While predicted miRNA target sites include many false positives [e.g. 78], and experimental work would be required to confirm a biological role, it is interesting to note that Kallithea Virus’ closest relative (Oryctes rhinoceros Nudivirus) does not encode a detectable homolog of the miRNA and contains only 4 predicted binding sites (miRanda score ≥150).

### Virus prevalence is high in laboratory stocks and cell culture

To detect the BLAST-candidate and siRNA-candidate viruses in previously published *Drosophila* studies, we mapped up to 2 million reads from each of 9656 publicly available fly and cell culture RNAseq and small-RNA datasets to the new and previously described *Drosophila* viruses. Around 33% of ‘run’ datasets contained viral reads above a threshold of 100 reads per million (rpm), representing 39% of ‘samples’ and 58% of submitted ‘projects’ (Figure 4, S23, S24). The proportion of positive samples varied with log-threshold, so that 53% had at least one virus at ≥10 rpm, but 17% of runs had at least one virus at ≥1000 rpm. These rates are slightly lower than, but not dissimilar to, previous estimates from serial passage of fly stocks, which found around 40% of fly stocks were infected [14]. BLAST-candidate and siRNA candidate viruses were both found, and by noting their co-occurrence we were able to identify several siRNA-candidates as component parts of other virus genomes (e.g. segments of Bloomfield Virus and the second segment of Craigie’s Hill Virus were initially identified as siRNA Candidate Viruses). The presence of BLAST-candidate and siRNA-candidate viruses in laboratory cultures supports these as *bona fide* infections of *Drosophila*, and demonstrates the utility of the viRNA signature as a marker for viruses.

**Figure 4:**
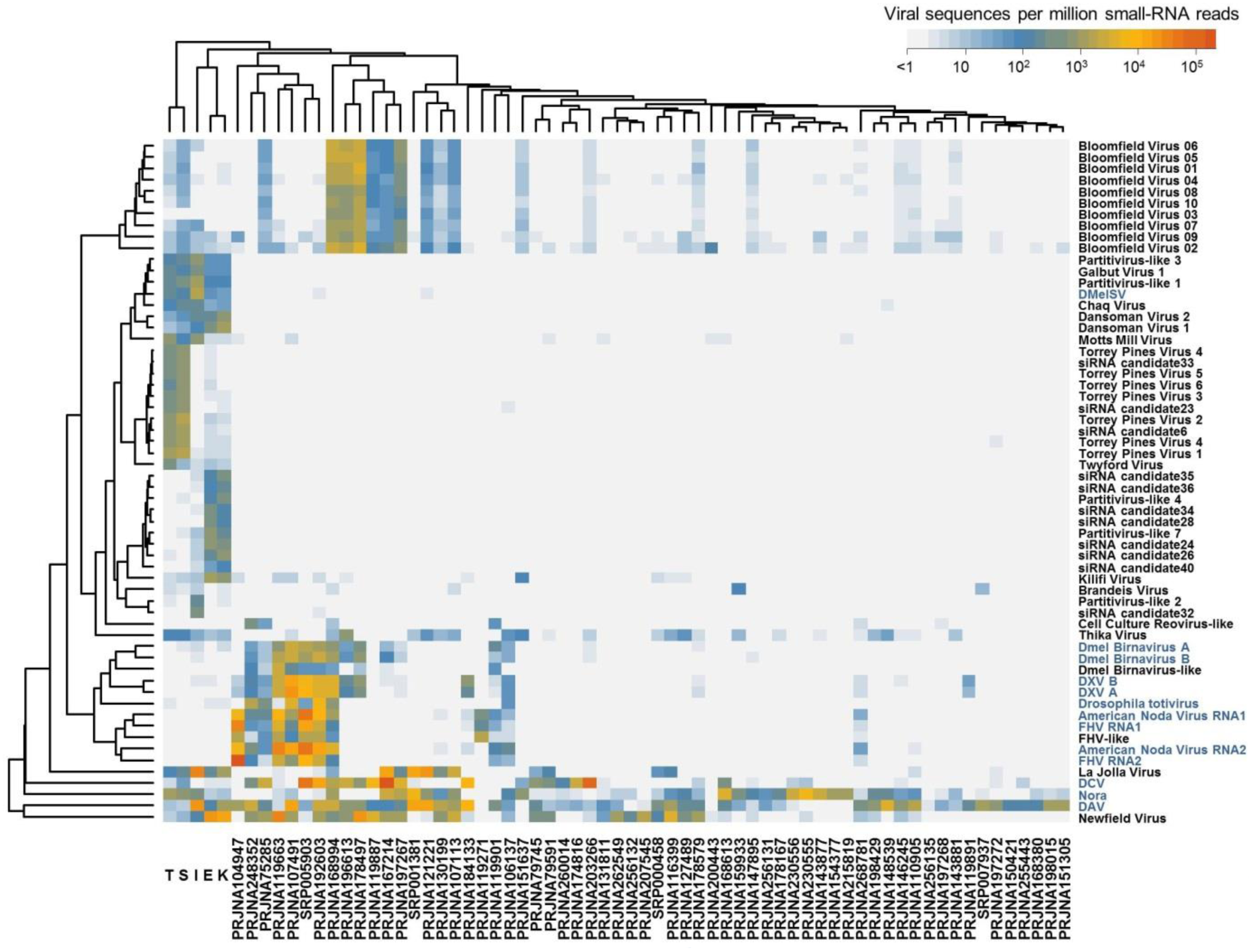
Virus-derived small RNA reads in publicly available RNA datasets. To discover the distribution of viruses in laboratory cultures (including cell culture) the first two million reads from each of 9656 publicly available RNAseq and siRNA datasets were mapped to all previously reported and newly discovered RNA viruses. The heatmap illustrates the virus community across 62 small RNA sequencing projects (averaging across read sets within projects) and the metagenomic discovery pools E, I, K, S, and T. Cells are coloured by the number of viRNAs per million total mappable reads, and rows and columns have been re-ordered to cluster similar viruses and similar samples. Only datasets and viruses with >100 rpm in at least one cell are included here. This analysis shows that many of the viruses, including newly discovered ones such as Newfield Virus, are extremely common in laboratory culture (including cell cultures) and demonstrates they are able to replicate in *Drosophila*. The analysis also suggests a clear difference in the virus community between the wild collected flies and laboratory flies. Previously described viruses are named in blue, new viruses and virus-like sequences in black. Data for all public RNA datasets are plotted S23 and summarized read counts for this figure in supporting information S21. A plot of exemplar cell culture datasets is given in supporting information S26 and raw counts from all public datasets are provided in supporting information S24.

Based on the 100 rpm threshold, DAV (1025 of 9656 datasets), DCV (979 datasets), Nora Virus (629 datasets), Newfield Virus (483 datasets), FHV, Drosophila Totivirus, American Nodavirus, Drosophila Birnavirus, DMelSV, Thika Virus, Kilifi Virus, La Jolla Virus, Craigie’s Hill Virus, Bloomfield Virus, Chaq Virus and Galbut Virus were all present in public datasets (Figure 4, S23, S24). However, some viruses (Kilifi Virus, Craigie’s Hill Virus, Chaq Virus, Galbut Virus, DMelSV) were extremely rare, appearing in 12 or fewer datasets. It is widely known that *Drosophila* cell culture harbours many viruses [21], and we identified multiple viruses and occasionally high numbers of viral reads in cell culture datasets. For example, reads mapping to nine different viruses were found in datasets SRR770283 and SRR770284 (Piwi CLIP-Seq in OSS cells [79]) and ≥70% of reads from SRR609669-SRR609671 were viral in origin (total RNA from piRNA-pathway knock-downs [80]). Supporting information S26 presents virus presence/absence for widely used *Drosophila* Cell cultures [81]. RNAseq datasets were also available for species other than *D. melanogaster*, and these too included virus-like sequences. We detected DAV in *D. pseudoobscura*, *D. virilis*, *D. bipectinata*, *D. ercepeae* and *D. willistoni*, DCV in *D. simulans*, *D. ananassae* and *D. mojavensis*, Nora Virus in *D. simulans*, *D. ananassae* and *D. mojavensis*, Thika Virus in *D. virilis* and *D. ficusphila*, Kilifi Virus in *D. bipectinata*, La Jolla Virus in *D. simulans*, and Bloomfield Virus in *D. virilis*.

Given the presence of viruses in public RNAseq and siRNA datasets, we selected a subset of 2188 datasets to perform viral discovery by *de novo* assembly. In adult *D. melanogaster* this identified a novel Picorna-like virus (present in 3 datasets among 9656) and a novel Negevirus (present in 10 datasets among 9656). We have provisionally named these Berkeley Virus and Brandeis Virus, respectively (S6, S25). The survey also identified a novel Totivirus and several (possibly fragmentary or non-coding) Reovirus segments in cell culture, at least one of which is widespread (214 datasets; Figure 4, S23, S24). The small number of novel viruses we identified in fly stocks over and above those described previously, may suggest that few further *Drosophila* viruses remain to be found regularly infecting laboratory stocks.

### *Drosophila* viruses are common and widely distributed in the field

To infer viral prevalence and distribution in wild flies we used RT-PCR to assay for the presence of 16 different viruses in a total of 1635 *D. melanogaster* and 658 *D. simulans* adults sampled from 17 locations across the world (Figure 5; S27). Excluding siRNA-candidate viruses, the most prevalent virus in large samples of *D. melanogaster* was La Jolla Virus (12/16 locations, 8.6% of flies averaged across locations) and the rarest was Twyford Virus (1/16 locations, average 0.3% of flies). Of those detected in large samples of *D. simulans*, the most prevalent was Thika Virus (4/7 locations, average 4.5% of flies) while the rarest was Motts Mill Virus (1/7 locations, average 0.2% of flies). Despite their high prevalence in the lab and presence in the metagenomic pools, DCV and Newfield Virus were not detected at any of the 17 locations, and Twyford Virus, DMelSV, Dansoman Virus and Craigie’s Hill virus were not detected in *D. simulans* (Figure 5; S27). Although sampling locations varied substantially in overall viral prevalence (in Athens GA >80% of *D. melanogaster* carried a virus; in Marrakesh less than 10%) there was no clear geographic structure in viral prevalence (Figure 5; S28), and for most viruses prevalence was not correlated between *D. melanogaster* and *D. simulans* from the same location. Excluding siRNA-candidate viruses, around 30% of *D. melanogaster* individuals and 13% of *D. simulans* individuals carried at least one virus, and over 6% of *D. melanogaster* individuals carried more than one virus (S29). We are unable to explain the unusually high viral prevalence in *D. melanogaster* sampled from Athens (GA, USA)—flies were separated to individual vials within a few hours, and *D. simulans* and *D. melanogaster* were netted together, but *D. simulans* did not show unusually high virus prevalence (S28).

**Figure 5:**
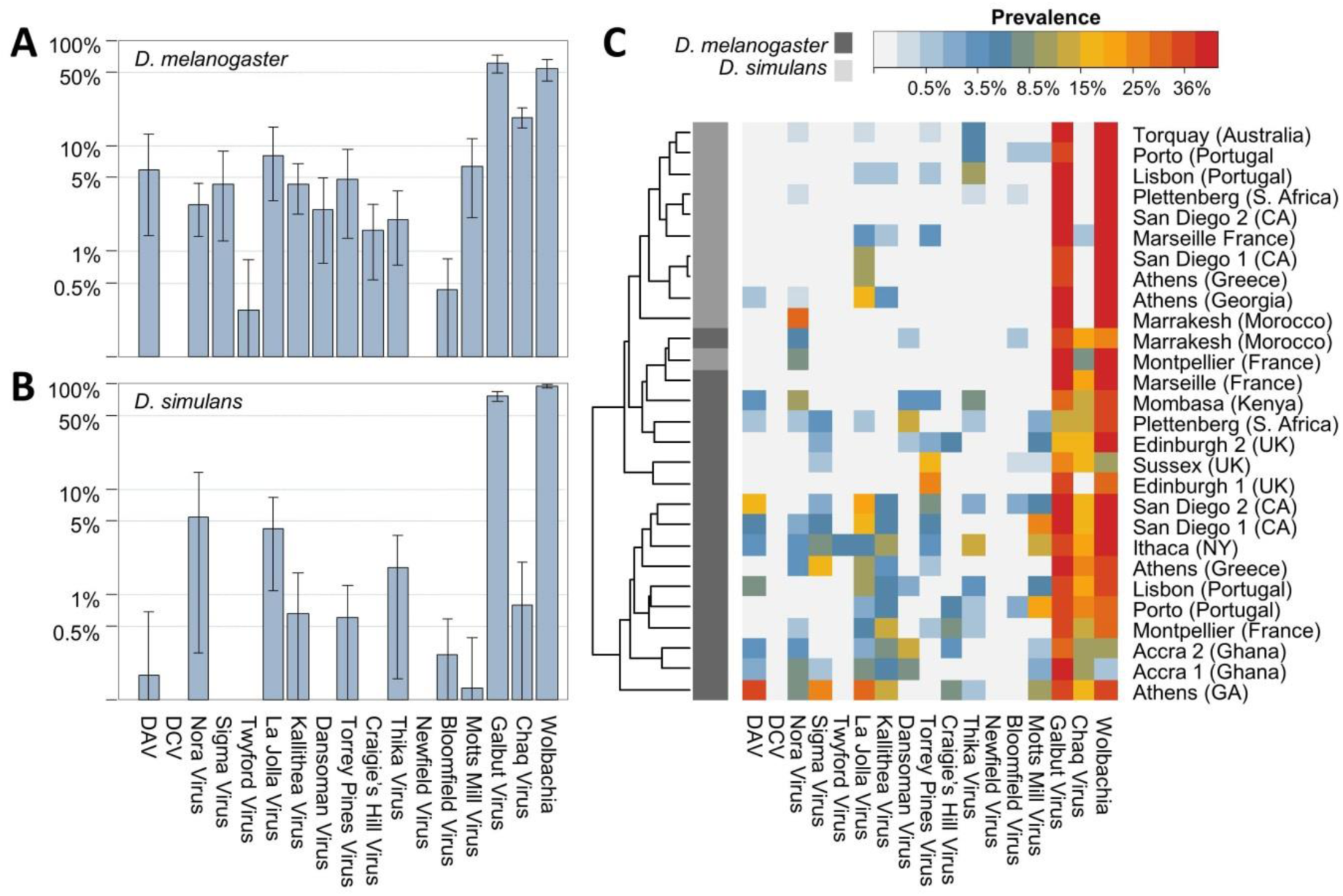
Distribution and prevalence of *Drosophila* viruses in the wild. Individuals and small bulks of adults were collected from 17 locations around the world and surveyed for the presence of viruses and *Wolbachia* by RT-PCR. Charts show mean global viral prevalence (averaged across populations) in (A) *D. melanogaster* and in (B) *D. simulans*, plotted on a log scale. Error bars represent 95% bootstrap intervals across populations. Panel (C) shows a heat-map of viral prevalence (locations by viruses), coloured on a log scale according to prevalence. Hosts are marked in dark grey (*D. melanogaster)* and light grey (*D. simulans*), and locations are given with collection year. Collections are clustered according to their similarity in virus prevalence. Supporting information S28 provides prevalence plots by location, and raw counts and location information are provided in supporting information S27.

In contrast to the other viruses, the novel siRNA-candidate viruses Galbut Virus and Chaq Virus often displayed extremely high prevalence. In *D. melanogaster*, Galbut Virus ranged from 13% in Plettenberg to 100% in Accra, and Chaq Virus from <5% in Edinburgh to 35% in Porto. In *D. simulans*, Galbut Virus ranged from 57% (Athens) to 76% (Torquay, Australia), although Chaq Virus was not highly prevalent (only 2/7 locations, at low prevalence). These rates are much higher than for the other viruses, and the inclusion of siRNA candidate viruses in overall infection rates brings many populations to ≥70% of flies carrying at least one virus (Figure 5, S28, S29). The high prevalence of these ‘siRNA-candidate’ viruses is surprising, and could perhaps imply an alternative (non-viral) origin for the sequences. However, we believe a viral origin is supported by a combination of the viral-like siRNA signature, their absence from *D. melanogaster* genomic reads, their close relationship to unclassified insect transcriptome sequences (Figure 1; S7), and their occasionally low prevalence (S28).

For DMelSV, which is the only virus previously surveyed on a large scale [reviewed in 15], our data agree closely with earlier estimates based on other assays: here 4.6% of *D. melanogaster* infected *versus* 2.8% in [22] and 5.0% in [23] (DMelSV is absent from *D. simulans*). Nevertheless, our estimates more generally should be treated with some caution as RT-PCR assays are unlikely to be reliable for all virus genotypes, leading to PCR failure for divergent haplotypes and thus potentially underestimation of prevalence.

Virus prevalence was strikingly different between publicly available RNA datasets and our field survey (Figure 4). For example, we identified Newfield Virus and DCV in many fly stocks and cell cultures but very rarely in the field (compare Figure 5 and S23; see also Figure 4), whereas Kallithea Virus and Motts Mill Virus were common in the wild (4.6% and 6.7% global average prevalence in *D. melanogaster*; Figure 5) but absent from DNA and RNA public datasets. The case of DCV is particularly striking as early surveys assayed by serial passage in laboratory cultures suggested DCV may be common in the field [14], as did PCR surveys of recently established laboratory stocks [82]. In the case of Newfield Virus and DCV, a downward collection bias, for example if high titre flies do not get collected, may explain the result. However, for viruses which have higher prevalence in the field than the lab such as Kallithea Virus and Motts Mill Virus, and also the siRNA candidates Galbut Virus and Chaq Virus, it seems likely that differences in (e.g.) transmission ecology between the lab and field may explain the disparity.

### *Wolbachia* prevalence is not detectably correlated with virus prevalence

Infection with the bacterial endosymbiont *Wolbachia pipientis* has previously been shown to confer protection against secondary infection by some RNA [9,10] but not DNA [83] viruses in insects. We therefore surveyed *Wolbachia* prevalence by PCR in the wild flies, and tested whether *Wolbachia* was correlated with virus presence. We found *Wolbachia* at detectable levels in all populations, ranging from 1.6% of individuals (Accra) to 98% (Edinburgh) in *D. melanogaster* and from 82% (Plettenberg) to 100% (Marseille) in *D. simulans* (Figure 5, S28). As expected [e.g. 84], overall *Wolbachia* prevalence was higher in *D. simulans* (∼90%) than in *D. melanogaster* (∼50%). We could not detect any consistent correlation across populations between the prevalence of *Wolbachia* and that of any virus (S30), nor could we detect any association between *Wolbachia* and viral infection status within populations (contingency table tests with *p*-values combined across populations using Fisher’s method; all *p*>0.05). While this may indicate that *Wolbachia*-mediated antiviral protection is not an important determinant of viral infection in the field, our relatively small sample sizes (median *n*=63 flies per location) mean that our power to detect an interaction is low. In addition it is unclear whether a positive or negative association should be expected, as some *Wolbachia*-host combinations appear to protect via increased tolerance rather than increased resistance [85], not all strains are highly protective [86], virus titre may correlate with *Wolbachia* titre even if there is no presence-absence relationship, and for at least one DNA virus resistance is decreased [83].

### Timescales of virus evolution

To provide a preliminary examination of the evolutionary and ecological dynamics of previously described and novel viruses we used a time-sampled Bayesian phylogenetic approach to infer dates of common ancestry and substitution rates [87,88]. To allow for differences in data coverage across (partial) viral genomes, and to allow for potential recombination, we divided alignments into blocks. These blocks were permitted to vary in mutation-rate (substitution-rate) and date parameters. For Nora Virus and DAV we were able to use early genomic samples [20,89] to aid time calibration, and we estimated the all-sites mutation rate for these viruses at approximately 4 to 8×10^-4^ nt^-1^ yr^-1^ (Nora Virus; range across blocks, posterior medians) and 2 to 6×10^-4^ nt^-1^ yr^-1^ (DAV). However, short sampling timescales means that these rates are estimated with low precision, resulting in a spread of 95% highest posterior density (HPD) credibility intervals across blocks of 2 to 12×10^-4^ nt^-1^ yr^-1^ and 1 to 9×10^-4^ nt^-1^ yr^-1^, respectively. This range falls within the expected range for single-stranded RNA viruses [90].

These mutation rates imply dates for the most recent common ancestors of DAV and Nora Virus (MRCA for known extant lineages) of approximately 100-200 years ago (range across blocks; range of credibility intervals across blocks 70-320 years) and 50-300 years ago (range across blocks; range of credibility intervals across blocks 33-630 years), respectively. This rapid movement of viruses is consistent with the rapid global movement that has been seen in transposable elements [91], *Wolbachia* [92], and even *Drosophila* genomes [93]. Nevertheless, inferred dates of a few hundred years in the past should be treated with caution, as weak constraint can lead to substantial underestimates of the true time to common ancestry [e.g. 94]. For the other nine RNA viruses analysed, sample sizes were smaller and dated samples were only available from a 5-year window, making unconstrained estimates of mutation rate and MRCA dates difficult so that quantitative estimates for these viruses should be treated with caution (results inferred using a strong prior on mutation-rate are presented in S31 and S33).

### Some viruses move repeatedly between *D. melanogaster* and *D. simulans*

Viruses detected in both *D. melanogaster* and *D. simulans* might either be reciprocally monophyletic, with substantial divergence between host-specific lineages, or the hosts may be distributed across the virus phylogeny, if between-host movement is common. By incorporating information on host species (*D. melanogaster vs. D. simulans*) in the phylogenetic analysis we were able detect past movement of RNA viruses between these host species (e.g. Figure 6) and obtain rough estimates of host switching rate [87,88] (S31 and S33). These rate estimates have very low precision, associated with the relatively broad posterior estimates for substitution rate, and should be treated with caution. Nevertheless, there appeared to be substantial variation in host-switching rate amongst the seven RNA viruses that were detected in both hosts, with Chaq Virus showing the lowest rate and La Jolla Virus the highest (Figure 6; supporting information S31 and S33). A similar analysis of inter-continental geographical movement suggests substantial genetic structuring between geographic regions may be present in some viruses (S31 and S33), although again further time-sampled data are required for the analyses to provide useful precision.

**Figure 6:**
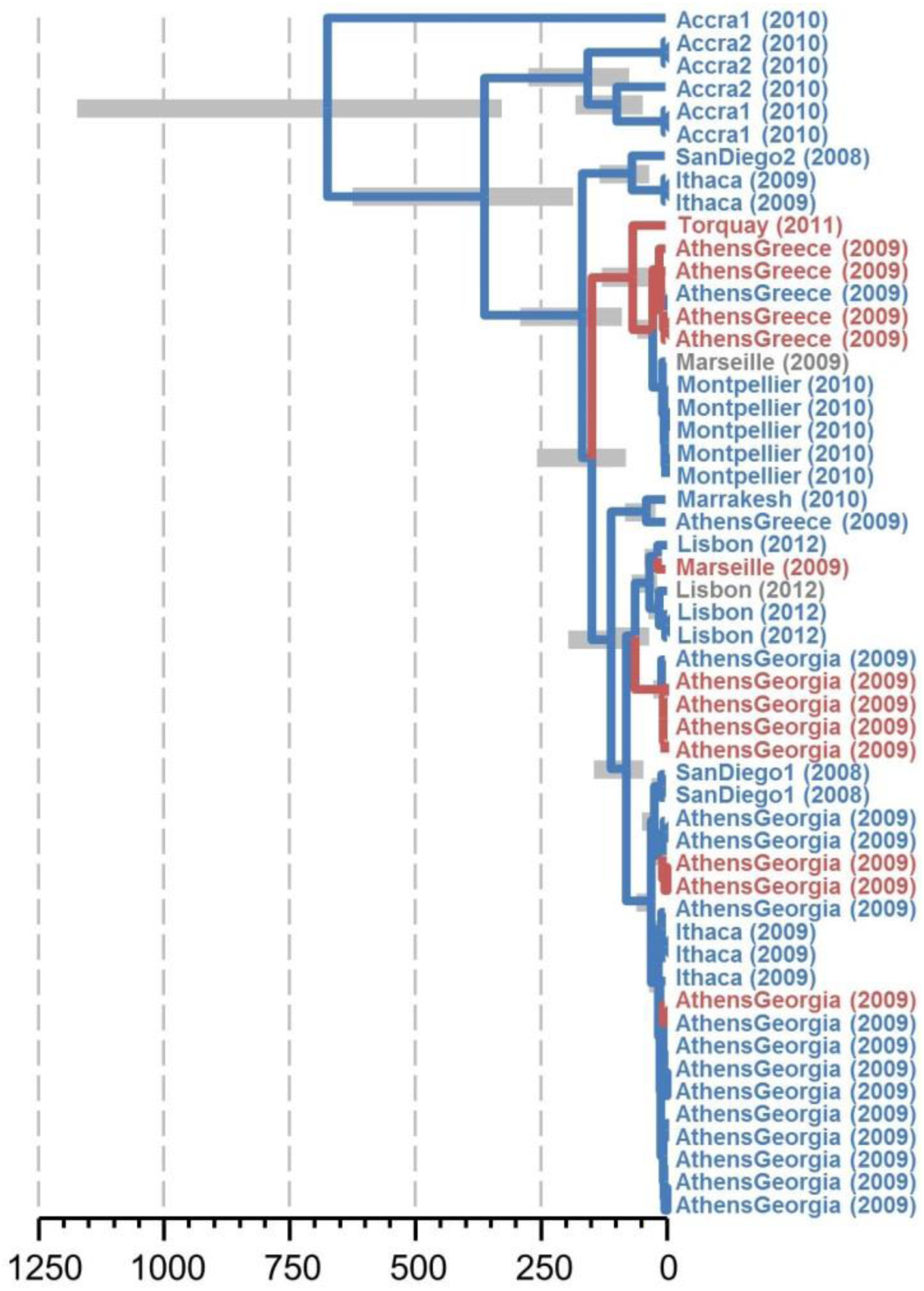
Host-switching and geographic structure in La Jolla Virus. The inferred hosts over time for a 700nt alignment block of La Jolla Virus. Location labels (and year of collection) are coloured by host species (*D. melanogaster* blue, *D. simulans* red) and branches are coloured by the inferred host, suggesting approximately seven host-switches in the tree. An estimated timescale is given in years before present, with 95% credibility intervals for node dates. However, the absolute timescale is tightly constrained by the prior assumption that RNA virus mutation rates fall between 1×10^-4^ and11×10^-4^ mutations per site per year, and given the limited temporal information in the data, should be treated with caution. Note that while sequences generally cluster by location, there is substantial mixing on a global scale. The underlying BEAST xml file (including the alignment and model specification), along with the resulting maximum clade credibility tree file and summaries of posterior distributions, are provided in supporting information S32 and S33.

### DAV and Nora Virus proteins are highly conserved with little evidence for positive selection

To quantify patterns of selection acting on protein-coding sequences, we applied a phylogenetic approach to infer relative rates of synonymous (*d*S) and non-synonymous (*d*N) substitution [see 95] in 11 of the viruses (quantified as *d*N-*d*S or *d*N/*d*S; S31, S34, S35). For most viruses, sample sizes were insufficient to obtain robust inferences of selection, however in general the protein sequences were strongly constrained (S31). DAV and Nora Virus, which had the most comprehensive sampling, showed among the lowest mean *d*N/*d*S ratios (0.25 and 0.24, respectively), although only Nora Virus displayed codons with strong evidence of positive selection. There were no clear patterns of *d*N/*d*S variation across genes within DAV or Nora virus genomes (S35), although 3 of the 4 positively selected codons were in the Nora Virus capsid [96], perhaps indicative of host-mediated selection. There was no evidence for positive selection or an elevated *d*N/*d*S ratio in the viral suppressor of RNAi encoded by Nora Virus [38], despite apparently rapid host-specialisation in this gene [12] and the rapid adaptive evolution seen in its antagonist, *Drosophila* Ago2 [47]. Nevertheless, within-species sampling may be unlikely to detect species-wide selective pressure [97] and the disparity in evolutionary timescales of host and virus may mean that even strong reciprocal arms races are not reflected in elevated viral *d*N/*d*S ratios (i.e. the time of common ancestry for the viruses is too recent for the host to have driven multiple substitutions within the sampling timeframe [98]).

## Conclusions

We have identified around 20 new viruses associated with *D. melanogaster* in the wild, and shown that some of them are common in the laboratory and the field. A substantial fraction of the virus lineages described here are newly (or only recently [33,57]) reported to infect arthropods. These include the novel ‘siRNA candidate’ viruses with uncertain affiliation, and new lineages related to Partitiviruses, Sobemoviruses and Poleroviruses, and Picornavirales. The presence of these viruses in cell culture and laboratory stocks, their absence from fly genomes, and the presence of virus-derived 21nt small RNAs all support the majority as *bona fide Drosophila* infections. *Drosophila*-associated viruses include many with positive sense RNA genomes (Dicistroviridae, Permutotetraviridae, Flaviviridae, Iflaviridae, Nodaviridae, Negeviruses, and others), negative sense RNA genomes (Rhabdoviridae; relatives of Bunyaviridae), and double-stranded RNA genomes (Birnaviridae, Totiviridae, Partitiviridae, Reoviridae), but few with DNA genomes (only the Kallithea and *Drosophila innubila* Nudiviruses [55] to date) and no retroviruses (*cf.* ‘Errantivirus’ endogenous retroviruses [99]; retro-elements or retrotransposons that are usually transmitted as genomic integrations). It therefore seems increasingly unlikely that the apparent wealth of RNA viruses and paucity of retroviruses and DNA viruses in *Drosophila* represents a sampling artefact, and this may instead reflect underlying *Drosophila* ecology or immune function.

This newly discovered diversity of *Drosophila* viruses will prove valuable for the use of *D. melanogaster* as a model for viral infection and antiviral immunity. First, these data will facilitate the curation and management of laboratory stocks and cell cultures. Second, they will facilitate the isolation and culture of viruses for future experimental work, including natural pathogens of *Drosophila* that are closely related to medically or economically important viruses such as Flaviviruses and Bunyaviruses, or Chronic Bee Paralysis Virus. Third, our analyses of virus prevalence, distribution, and dynamics will provide an informed evolutionary and ecological context to develop *D. melanogaster* and its relatives as a model system for viral epidemiology and host-switching. In combination, we hope this work will provide a key reference point for future studies of *Drosophila*-virus interaction.

## Materials and Methods

### Sample collections

For metagenomic viral discovery, five large collections of *Drosophila melanogaster* and other sympatric drosophilid species were made in 2010 by netting flies from fruit bait. For UK samples (denoted S and T), flies were morphologically screened to exclude species other than *D. melanogaster*. For non-UK samples (denoted E, I and K), flies were screened to bear a superficial similarity (small, pale, stripes) to *D. melanogaster*. Samples E and K, each of *ca.* 700 adult pale-bodied Drosophilidae were collected in Kilifi (Kenya) by JA in July 2010. Sample I of *ca.* 650 adult pale-bodied Drosophilidae were collected in Ithaca (NY, USA) by BPL in August 2010. Sample ‘S’ of *ca.* 1250 adult *D. melanogaster* were collected in Sussex (UK) by DJO and EHB in August 2010. Sample ‘T’ of *ca.* 1700 adult *D. melanogaster* were collected in Twyford (UK) by DJO and Alasdair Hood in July 2010. In each case flies were maintained in groups of up to 200 for 5-10 days on a hard-agar/sugar medium before maceration under liquid nitrogen. RNA extraction was performed separately on each of the five samples.

For analyses of prevalence and sequence evolution, collections of *D. melanogaster* and *D. simulans* were made at the following locations between November 2008 and October 2012: Thika (Kenya) by John Pool in January 2009; San Diego (CA, USA) by DJO in November 2008; Edinburgh (UK) by DJO in August 2009; Athens (Greece) by Natasa Fytrou in June 2009; Marseille (France) by Nicolas Gompel in July 2009; Ithaca (NY, USA) by BPL in September 2009; Athens (GA, USA) by PRH in September 2009; Sussex (UK) by DJO in August 2009; Two locations in Accra (Ghana) by CLW and JA in January 2010; Plettenburg (South Africa) by Francis Jiggins in January 2010; Edinburgh (UK) by CLW in September 2009; Marrakech (Morocco) by CLW in September 2010; Montpellier (France) by PRH in September 2010; Torquay (Australia) by BL in February 2011; Lisbon (Portugal) by DJO in October 2012; Porto (Portugal) by DJO in October 2012 (S27 for details). In each case flies were aspirated or netted from fruit bait at intervals of 24 hours or less, and maintained individually in isolation for up to 10 days on a sugar/hard-agar medium prior to freezing at - 80°C. In addition, a small number of laboratory-stock flies were provided by David Finnegan (University of Edinburgh) in February 2008 and February 2010.

### Metagenomic and small-RNA sequencing

All raw reads have been submitted to the Sequence Read Archive under project accession SRP056120. RNA was extracted using Trizol® (Life Technologies) and DNAse treated (Life Technologies) according to the manufacturer’s instructions. For metagenomic RNAseq and small-RNA sequencing, aliquots of the samples E, I, K, S, and T were initially mixed into two pools: Dmel/UK (denoted ST) and mixed-species/non-UK (EIK). RNAseq was performed by Edinburgh Genomics (Edinburgh). Following ribosome-depletion using two cycles of RiboMinus^TM^ (Life Technologies), around 40 million high-quality paired-end 100nt Illumina reads were generated from a single sequencing lane of each library (accessions SRR1914484 and SRR1914527). Small-RNA sequencing from the same two pools was performed with a periodate oxidation step to reduce the relative ligation efficiency of miRNAs, which do not carry 3'-Ribose 2'O-methylation [e.g. 76], and sequenced by Edinburgh Genomics (Edinburgh). An equivalent library, without periodate oxidation, was constructed and sequenced by BGI (Hong Kong) These libraries generated between 12 and 20 million small-RNA reads each (SRR1914671, SRR1914716, SRR1914775, SRR1914792). After preliminary analysis, RNA aliquots of all five pools were mixed (mix denoted EIKST) and RNAseq was repeated by BGI (Hong Kong) following either double-stranded nuclease normalization (around 65 million high-quality paired-end 90nt Illumina reads; SRR1914412), or ribosome depletion using Ribo-Zero (Illumina; around 65 million high-quality paired-end 90nt Illumina reads; SRR1914445). Finally, individual small-RNA libraries were constructed for each of the five samples E, I, K, S, and T, without periodate oxidation using both NEBnext® adaptors (New England Biolabs) (datasets SRR1914946, SRR1914952, SRR1914955, SRR1914957, SRR1914959), and equivalent ‘High Definition’ adaptors (datasets SRR1914958, SRR1914956, SRR1914954, SRR1914948, SRR1914945), which seek to reduce ligation bias by incorporating random tetramers into the ligation adaptor [66]. These ten libraries were sequenced by Edinburgh Genomics (Edinburgh). Any small RNAs produced by synthesis (e.g. nematode 22G small RNAs) carry a 5' triphosphate, and are unlikely to be ligated by this protocol.

To estimate species-composition of the UK (ST) and non-UK (EIK) pools, RNAseq datasets were quality trimmed using ConDeTri2.0 and mapped using Bowtie2 [100] to a 359nt fragment of COI (position 1781-2139 in *D. melanogaster* reference mtDNA) drawn from multiple species. This identified the EIK RNA sample to have been a mixture of *D. simulans* (0.8%), *D. hamatofila* (1.2%), *D. ananassae* (12.4%), *Scaptodrosophila latifasciaeformis* (16%), *D. melanogaster* (24.1%), and *D. malerkotliana* (45.5%). It also identified a small amount of cross-species contamination in the *D. melanogaster* ST sample (0.4% *D. simulans*, 0.4% *D. immigrans*). To estimate the contribution of each of several potential sources of RNA, reads were also mapped to known *Drosophila* viruses, the *D. melanogaster, D simulans* and *D ananassae* genomes, *Wolbachia* from *D. melanogaster*, *D. simulans*, *D. willistoni*, *D. suzukii*, and *D. ananassae*, and the bacteria *Pseudomonas entomophila, Providencia sneebia, P. rettgeri, P. burhodogranariea, P. alcalifaciens, Gluconobacter morbifer, Enterococcus faecalis, Commensalibacter intestini, Acetobacter pomorum, A. tropicalis*, and *A. malorum* (S2). The percentage of mapping reads was quantified relative to total high quality reads.

### Identification of virus-like contigs by BLAST

All paired-end RNAseq datasets were combined and *de novo* assembled using three different approaches. First, all reads were assembled using Trinity (r2013-02-25; [50]). Second, data were digitally normalised using the “normalize-by-median-pct.py” script from the Khmer package, and then assembled using Trinity. Finally, this normalised dataset was also assembled using the Oases/Velvet pipeline (version. 0.2.08; [51]). Contigs are provided in supporting information S5. Using the longest inferred contig for each putative locus and a minimum contig length of 200nt, contigs were compared to the Genbank protein reference sequence database using BLASTx [52] with default search parameters and an e-value threshold of 1×10^-10^ to identify the top 10 hits. Contigs with no BLAST hit at this threshold were translated to identify the longest open reading frame in the contig, and the resulting amino acid sequence compared to Genbank protein reference sequence using BLASTp with default parameters and an e-value threshold of 1×10^-5^ to identify the top 10 hits. For each contig with BLAST hits, the phylogenetic hierarchy of the best hits was traversed upward to identify the lowest taxonomic classification displaying a 75% majority taxonomic agreement, and this was used to guide subsequent cross-assembly and manual curation using the SeqManPro assembler (DNAstar). The resulting BLAST-candidate virus sequences were then used as targets for (RT-)PCR and Sanger sequencing confirmation (both on discovery samples EIKST, and on individual flies), with primer design guided by assuming synteny with close relatives. Final sequences for phylogenetic analysis and submission to NCBI were generated either by PCR and Sanger sequencing, or from a majority consensus derived by remapping RNA-seq reads to the combined *de novo* / PCR contig (sequences have been submitted to Genbank as KP714070-KP714108 and KP757922-KP757936).

### Identification of virus-like contigs by small-RNA profile

The *Drosophila* Dcr2-Ago2 RNAi pathway processes double-stranded RNA derived from viruses and transposable elements into 21nt siRNAs [e.g. 7,8,101]. In the case of viruses, Dcr2 may process both replicative intermediates and fold-back structures. However the presence of replicative strand reads (i.e. negative strand reads in positive sense RNA viruses) demonstrates at that replicative intermediates are a major substrate for Dcr2. Thus, the presence of virus-derived 21nt siRNA sequences from both strands is diagnostic of an antiviral response against replicating viruses. The piRNA pathway also generates small RNAs (25-29nt piRNAs from transposable elements [67]), however, although viral piRNAs are found in *Drosophila* OSS cell culture expressing Piwi [17], these have not been detected in whole flies. Therefore the presence of large numbers of 21nt small RNAs, relative to total RNA or to 25-29nt small RNAs, provides corroborating evidence that virus-like sequences are recognised by the *Drosophila* immune system.

In addition, we reasoned that this signature of 21nt siRNAs could also be used to identify viral sequences amongst the substantial fraction of contigs that displayed no BLAST similarity to any sequence in the NCBI reference protein database. To quantify the siRNA profile of the *de novo* contigs we mapped the 19-29nt small RNAs from the ST and EIK datasets to all *de novo* contigs, recording all potential mapping locations. As a proxy for total RNA, we also re-mapped combined EIKST RNAseq data using the dataset generated by BGI (Hong Kong) with Ribo-Zero rRNA depletion. For each contig this resulted in a count per unit length of the number of 21-23nt siRNAs, the number of 25-29nt putative piRNAs, and the number of RNAseq reads. Using the well-studied *Drosophila* viruses DCV, DAV, Nora virus, and DMelSV as threshold a guide, we then used a high ratio of siRNA:RNAseq reads and a high ratio of siRNA:piRNA reads to corroborate the BLAST-candidate viruses outlined above, and to propose novel siRNA-candidate viruses amongst contigs lacking BLAST hits (Figure 3). Final siRNA-candidate sequences for phylogenetic analysis and submission to Genbank were generated either by RT-PCR and Sanger sequencing, or from a majority consensus derived by re-mapping RNA-seq reads to the each contig (Submitted to Genbank as KP757937-KP757993).

### Viral phylogenetic analyses

We used translations from the novel viruses to perform a BLASTp similarity search of the Genbank protein reference database to identify relatives and suitably conserved genomic fragments for phylogenetic inference. In addition, we searched the Genbank transcriptome shotgun archive (TSA: database tsa_nt) using tBLASTn [52] to identify potential virus sequences (i.e. sequences currently unannotated as viral) in recently generated high-throughput transcriptomes. Protein sequences from known viruses, novel viruses, and TSA candidate viruses were then aligned using T-Coffee ‘psicoffee’ [102], and some poorly aligned regions at the 5' and 3' ends of the selected fragment were manually identified and trimmed. Phylogenetic trees were inferred using MrBayes [103] under a protein substitution model that allowed model jumping between widely advocated amino acid substitution models, and gamma-distributed rate variation among sites. Two independent MCMC chains were run sampling every 100^th^ step until the posterior sample of tree topologies reached stationarity (i.e. standard deviation in split frequencies between chains dropping to <0.01) and the effective sample size of all parameters was >300 (after 25% burn-in). Maximum clade-credibility consensus trees were prepared by combining both MrBayes chains using TreeAnnotator from the BEAST package [88]. For most of the novel RNA viruses the phylogeny was inferred from the RNA polymerase, which tends to be highly conserved, but for Kallithea Virus (a DNA Nudivirus) we selected five loci that have previously been used in Nudivirus phylogenetics [104] and the phylogeny was inferred from their concatenated alignments.

### Properties of the siRNA profile and novel viral miRNAs

To quantify the relative number of small RNAs produced from each virus, RNAseq reads and small RNAs from the EIK and ST pools were re-mapped using Bowtie2 [100]. To quantify the length distribution and 5' base-composition, the small RNAs from 14 different sequencing runs were mapped to known miRNAs (mirBase [105]), known transposable elements (Flybase [106]), all viral genomes, and other parts of the *D. melanogaster* reference genome (Flybase). The 14 siRNA datasets comprised: each of E, I, K, S and T ligated using the NEBnext protocol according to manufacturer’s instructions; each ligated using the NEBnext protocol replacing oligos with ‘High Definition’ equivalents that incorporate four additional random bases [66]; and the EIK and ST pools sequenced by Edinburgh Genomics with and without periodate oxidation (BGI) to reduce miRNA ligation efficiency [e.g. 76]. To identify potential miRNA sequences in the large dsDNA genome of Kallithea Virus, all small-RNAs were combined and mapped to the Kallithea Virus genome using Bowtie2, and novel miRNAs were inferred from read numbers and predicted hairpin-folding using miRDeep2 with default parameters [75].

### Identification of virus-like sequences in public RNAseq and genomic datasets

Several viruses are known to be endemic in laboratory stocks and cell cultures of *D. melanogaster* [e.g. 14,21], and we expected that many of the novel viruses identified here may also be detectable in those datasets. We therefore searched the European Nucleotide Archive for Illumina, SOLiD and Roche454 sequencing ‘run’ datasets that derive from the Drosophilidae and subordinate taxa. This resulted in 14123 DNA sequencing ‘run’ datasets and 9656 RNAseq and siRNA sequencing ‘run’ datasets (as of 9^th^ May 2015) from each of which we were able to download and map the first 2 million forward-orientation reads to new and previously known RNA viruses using Bowtie2 ([100]; Bowtie for SOLiD colour-space reads). To allow for mismapping, an arbitrary threshold of 100 reads per million (rpm) was chosen as a detection limit for viruses to be recorded as present. This threshold suggests at least one virus was present in 33% of runs, but the proportion of positive samples decreased with increasing log(threshold), suggesting that true infection rates may be higher. Across the RNAseq and siRNA datasets the co-occurrence of BLAST-candidate viruses with siRNA-candidate viruses allowed us to tentatively assign some of the siRNA-candidate viruses as components of BLAST-candidate virus genomes (e.g. Bloomfield Virus and Craigie’s Hill Virus).

To detect additional novel RNA viruses present in these public datasets, but absent from the literature or from our metagenomic sequencing of wild flies, we further selected 2188 Illumina RNAseq datasets for viral discovery. The first 20 million read pairs (or as many as were available, if fewer) from each dataset were mapped to the *D. melanogaster* genome (Bowtie2, local alignment), and unmapped reads digitally normalised (Khmer) and *de novo* assembled using Trinity [50] as above. Resulting contigs of >3.5 kbp in length were compared to known viral proteins using BLASTx, and the top hits manually curated to create a second group of BLAST-candidate viruses (such sequences cannot be accepted by Genbank, and are instead provided in Supporting File S25).

We reasoned that the absence of virus-like sequences from *D. melanogaster* genomes can provide evidence that these sequences do not represent ‘fossilised’ endogenous viral elements (EVEs). However, if only a minority of individuals carry a particular EVE, and if genomes are derived primarily by mapping to a reference, the EVEs may not appear in assembled datasets. We therefore mapped all reads (forward and reverse reads mapped separately) from 779 *Drosophila* NEXUS genomic read datasets (representing 527 *D. melanogaster* genomes [60]) to known *Drosophila* viruses, BLAST-candidate viruses, and siRNA-candidate viruses. Because a small number of reads may mismap to viral sequences, potentially leading to false inference of an EVE, read numbers were quantified relative to target sequence length, and normalised to a 7kbp single-copy region of the *D. melanogaster* genome (as a segregating EVE present in a genome should generate reads at a rate approaching that of other loci in that genome).

### Population prevalence

We used RT-PCR to survey wild populations for the presence of four previously published *Drosophila* viruses (DCV, DAV, DMelSV and Nora Virus), ten novel BLAST-candidate viruses (Bloomfield Virus, Craigie's Hill Virus, Dansoman Virus, Kallithea Virus, La Jolla Virus, Motts Mill Virus, Newfield Virus, Thika Virus, Torrey Pines Virus, Twyford Virus) and two novel siRNA-candidate viruses (Chaq Virus and Galbut Virus). In addition, we surveyed the same RNA extractions by RT-PCR for the presence (but not titre) of *Wolbachia pipientis*, using primers for the *Wolbachia* surface protein (wsp_F1 GTCCAATARSTGATGARGAAAC and wsp_R1 CYGCACCAAYAGYRCTRTAAA).

This survey comprised a total of 1635 *D. melanogaster* individuals and 658 *D. simulans* individuals sampled across 17 locations (see ‘Sample collections’ above), with between 2 and 236 individuals assayed for each species at each location (median *n*=63). RNA extractions and random hexamer reverse transcription was performed on single flies (males and females) or small bulks of 2-20 flies (usually ≤10 males, with species initially assigned based on morphology). Fly species was confirmed using a competitive 3-primer PCR for the single-copy nuclear gene *Ago2*, which gives a 300 bp product in *D. simulans* and a 400bp product in *D. melanogaster* (Primers: Mel_SimF CCCTAAACCGCAAGGATGGAG, Dsim303R GTCCACTTGTGCGCCACATT, Dmel394R CCTTGCCTTGGGATATTATTAGGTT)

For each of these 16 viruses we designed up to four PCR assays based on conserved positions in the metagenomic RNAseq datasets, and these were tested on the E, I, K, S and T metagenomic discovery RNA pools and on a selected subset of wild-collected individual flies. Those primer sets that consistently gave repeatable bands in agreement with other assays for the same virus, and which could be sequenced to confirm band identity, were selected for use as assays for the wider geographic sampling (some designed as multiplex assays; details of primers and conditions are given in supporting information S36). To allow for the possibility of PCR failure caused by primer-site variants not detected in virus discovery or preliminary sequencing, we designated flies as nominally positive for a virus if at least one assay for that virus resulted in an RT-PCR product. Nevertheless, the potential for unknown sequence variants to prevent PCR amplification means that our estimates of viral prevalence could be considered lower bounds.

As only one fly needs to carry a virus for the bulk to test positive for that virus, the proportion of ‘positive’ single-fly and/or small-bulk assays does not directly estimate to the frequency of virus carriers. We therefore used a maximum likelihood approach to infer the underlying prevalence (and its log-likelihood confidence intervals). Briefly, we assumed that for each species/location combination there was a single underlying prevalence (i.e. fraction of flies carrying detectable virus) and that the number of flies testing positive or negative by PCR assay would be binomially distributed given this prevalence. Then, given the number of flies included in each small bulk, and the number of small bulks or single flies testing positive, we searched across the range of possible prevalence values (zero to one) to identify the prevalence which maximised the likelihood of the observed set of positive/negative PCR assays. It is possible that differences in the sex ratio between individual-fly assays (majority female) and small-bulk assays (majority male) could make this approach misleading if prevalence differs between males and females. However, based on likelihood ratio tests (identical prevalence *versus* small bulks differ from single flies) we were unable to identify any widespread significant differences between the sampling regimes. Specifically 4.4% of tests were nominally ‘significant’ at *α*=5% (as expected), and only two were ‘significant’ using the Benjamini-Hochberg approach to limit the false discovery rate to 5% (Galbut Virus in Accra-1, *Wolbachia* in Plettenberg).

### Viral phylodynamics and patterns of sequence evolution

Eleven of the viruses with high prevalence (DAV, Nora Virus, DMelSV, Craigie’s Hill Virus, Dansoman Virus, La Jolla Virus, Motts Mill Virus, Thika Virus, Torrey Pines Virus, Chaq Virus, and Galbut Virus) were selected for further sequence analyses. PCR products amplified during the prevalence survey were supplemented with additional amplicons and sequenced in both directions using BigDye V3.1 (Life Technologies). Reads were assembled into partial or near-complete genomes using SeqManPro (DNAstar) and all variable sites within or between chromatograms were examined by eye. Virus sequences that displayed substantial within-sample (within-fly or small bulk) heterozygosity for any amplicon, and therefore deemed likely to represent mixed infections, were discarded. Sequences were submitted to Genbank under accessions KP969454-KP970120.

Sequence alignments were divided into blocks, minimising missing data within blocks, and these alignment blocks were tested for evidence of recombination using GARD [107] from the HyPhy package under a GTR model of nucleotide substitution with inferred base frequencies and 4-category gamma-distributed rate variation between sites. Those blocks which displayed any evidence of recombination were further subdivided at the inferred recombination break points, to result in multiple alignment blocks for each virus.

Alignment blocks (containing no detectable recombination) were subjected to phylogenetic analysis using BEAST [88] under a strict-clock model with HKY substitution using inferred base frequencies, and 4-category gamma-distributed rate variation between sites. Separate substitution models with relative clock rates were used for 1^st^, 2^nd^, and 3^rd^ codon and for non-coding positions, with substitution models linked across alignment blocks. Time calibration information was incorporated using a uniform step prior on tip sample-dates based on the value and precision of sample freezing date, and molecular clock rate was unlinked between alignment blocks. As the limited sampling timespan meant that little rate information was available for most viruses, we chose to implement a strong prior on nucleotide clock-rate, based on published virus estimates [90]. For RNA viruses this was a log-normal distribution with mean 4×10^-4^ site^-1^ year^-1^ and 95% of the prior density between 1×10^-4^ and 11×10^-4^. Where greater time-sampling depth was available (DAV and Nora Virus) we ran models using a hierarchical prior for global nucleotide clock rate, in which clock rates for individual segments are drawn from a shared log-normal distribution. To infer rates of host-switching between *D. melanogaster* and *D simulans*, and rates of movement among broadly-defined sampling locations (Europe and North Africa, Sub Saharan Africa, North America, Australia, Laboratory Stocks), tips were labelled by location and host species and treated as discrete traits [e.g. 87,108]. All analyses assumed a standard neutral coalescent in a constant-size population, and other parameters took default values and priors. Two analyses were run for each model for up to 10^8^ steps sampling every 50 thousand steps, and convergence was inferred for all parameter values both visually from stationary distributions, and from effective sample sizes of >1000. In one case (Thika virus) time and date parameters mixed poorly, and the analyses were re-run using a hard upper limit to the nucleotide clock rate for one alignment block (2.5×10^-3^ site^-1^ yr^-1^). Posterior parameter estimates and maximum clade credibility trees were inferred from the two independent parallel runs combined after discarding 10% of steps of each as burn-in.

Patterns of synonymous (*dS*) and nonsynonymous (*dN*) substitution, and evidence for positive selection (*dN*-*dS*>0) were inferred using the FUBAR [95] analysis from the HyPhy package. The in-frame viral coding sequence alignment blocks (as used above in phylogenetic reconstruction) were analysed based on maximum clade-credibility tree topologies inferred using BEAST, with substitution parameters and branch lengths inferred by HyPhy. FUBAR provides a fast approximate approach to infer *dN-dS* and sites under selection by pre-computing a grid of conditional likelihoods for *dS* and *dN* (where the expectation of *dS* is 1) and reusing the precomputed values to infer the posterior distribution of *dN-dS* without recalculating likelihoods. The analysis used a 20 × 20 grid with 10 independent MCMC chains each providing 1000 subsamples from the posterior (each 5 × 10^8^ steps after 5 × 10^8^ burn-in steps). Sites were considered to display evidence of positive selection if more than 95% of the posterior sample for that site supported *dN*>*dS.*

## Author Contributions

CLW performed sample preparation, assays and sequencing with support from FMW, SR, DC, GF and J-MB. CLW, JFQ, AHB and EHB prepared small RNA libraries. FMW performed qPCR. CLW, PRH, JA, BPL, EHB, BL and DJO performed field collections. DJO conceived and designed the study and performed all analyses. DJO, FMW, AHB and BPL wrote the paper, with contributions from other authors. We thank four anonymous reviewers and the editor for their thoughtful comments on the manuscript.

## Acknowledgements

We thank David Finnegan, Natasa Fytrou, Rosina Kyerematen, Charles Mbogo and the Kenya Medical Research Institute, Jonathan Barber, Sergio Castrezana, Kelly Dyer, Alison Duncan, Susana Gomes dos Santos Barber, Nicolas Gompel, Daniel Halligan, Dennis Hartley, Alasdair Hood, Therese Markow, Keith & Sue Obbard, John Pool, Carolyn Riddell and John Sinclair for providing flies, permission to collect flies, and/or logistical support in the field. We thank John Davey, Urmi Trivedi, Sujai Kumar, Timothee Cezard and members of Edinburgh Genomics for help and advice with sequence analyses. We thank Gytis Dudas, Francis Jiggins, Andrew Rambaut, Ronald van Rij, Pedro Vale, Lena Bayer-Wilfert, and members of the Obbard and Vale labs for discussion.

## Financial Disclosure

This work was funded by a Wellcome Trust Research Career Development Fellowship (WT085064) to DJO, and work in DJO’s and AHB’s labs is supported by a Wellcome Trust strategic award to the Centre for Immunity, Infection and Evolution (WT095831). PRH was supported by a fellowship from the UK Natural Environment Research Council (NE/G013195/1). The funders had no role in study design, data collection and analysis, decision to publish, or preparation of the manuscript.

## Supporting Information

### S1 Mappable reads

The proportion of RNAseq and small (17-29nt) reads that map to characterised *D. melanogaster* transposable elements, *Drosophila* miRNAs, the remainder of the *D. melanogaster* genome, *Wolbachia* and other bacteria, the four ‘classical’ viruses (DAV, DCV, DMelSV and Nora Virus), the new named viruses, other BLAST-candidate viruses, and siRNA-candidate viruses. Counts exclude unmapped reads, and reads mapping to *Drosophila* ribosomal sequences. EIKST refers to the metagenomic pools or mixtures thereof, ‘BGI’ and ‘EG’ indicate sequencing was performed by the Bejing Genomics Institute or Edinburgh Genomics (respectively), ‘HD’ indicates the use of ‘High Definition’ ligation adapters for small RNA sequencing, ‘RM’ and ‘RZ’ indicate to use of rRNA depletion by RiboMinus or Ribo-Zero, and ‘DSN’ indicates double-stranded nuclease normalisation. Raw counts data are provided in S2.

### S2 Read count data for S1

Counts of raw metagenomic reads mapping to *Drosophila*, bacteria, and viruses presented in supporting information S1.

### S3 qRT-PCR of metagenomic virus pools

RNA quantification for DCV, DAV, Nora Virus, DMelSV, Thika Virus, Motts Mill Virus, Torrey Pines Virus, Newfield Virus, Twyford Virus, La Jolla Virus and Craigie’s Hill Virus present in each of the five metagenomic pools E, I, K, S, and T, quantified by qRT-PCR relative to the *Drosophila* ribosomal protein gene RpL32. Note that DCV and Twyford Virus were each detectable in a single pool. Error bars represent 80% credibility intervals for means, based on two or three replicates per sample and assuming the efficiencies inferred from dilution series (S36). Virus presence/absence agrees well with small RNA data (S1) but qRT-PCR quantification may be unreliable given the high sequence diversity in these virus pools. Raw CT data are given in S4.

### S4 qRT-PCR CT values for S3

Raw qRT-PCR CT values for the analysis presented in supporting information S3

### S5 Metagenomic contigs

Raw metagenomic contigs derived from Trinity [50], Trinity with normalised data, and Oases [51] with normalised data, are provided in fasta format. The majority of contigs are likely to derive from *Drosophila* and associated microbiota. The contigs are uncurated, and are likely to include chimeric assemblies. As such, they are not suitable for submission to Genbank, and should be treated with caution.

### S6 Detailed description of new viruses

Detailed descriptions of BLAST-candidate virus fragments and named siRNA-candidate viruses

### S7 Phylogenetic trees

Mid-point rooted maximum clade-credibility trees showing the inferred relationship between new and previously known *Drosophila*–associated viruses (red), previously published viruses from other taxa (black) and sequences from the Transcriptome Shotgun Archive (blue). Bayesian posterior support is shown for second-order nodes and above and where space permits, and the scale is given in amino-acid substitutions per site. Genbank nucleic acid or protein identifiers are given after each sequence. (**A**) Craigie’s Hill Virus and Nodaviruses; (**B**) Kilifi Virus, Thika Virus, DCV, and Dicistroviruses; (**C**) Bloomfield Virus, Torrey Pines Virus, and Reoviruses; (**D**) Newfield Virus, DAV, and Permutotetraviruses; (**E**) Galbut Virus and TSA sequences; (**F**) La Jolla Virus, Twyford Virus, and a closely related Iflaviruses; (**G**) Motts Mill Virus, Sobemoviruses and Poleroviruses; (**H**) Charvil Virus and Flaviviruses; (**I**) Dansoman Virus and viruses related to Chronic Bee Paralysis Virus; (**J**) Chaq Virus and TSA sequences; (**K**) Brandeis Virus and related Negeviruses; (**L**) Berkeley Virus, DCV and a selection of Picornavirales sequences, (**M**) Partitiviruses; (**N**) Bunyaviruses; (O) Kallithea Virus and Nudiviruses (Note that *D. innubila* Nudivirus [55] is excluded because the relevant loci are unavailable). Alignments are provided in supporting information S8, and maximum clade credibility trees are provided in supporting information S9.

### S8 Alignments for Figure 1 and S7

Protein sequence alignments used to generate the trees presented in Figure 1 and supporting information S7 are provided in nexus format, along with the MrBayes commands used to perform the analyses.

### S9 Tree files for Figure 1 and S7

Maximum clade credibility trees with Bayesian posterior support are provided in BEAST nexus format, suitable for display with FigTree.

### S10 Read count data for Figure 2

Raw counts of 17-29nt small RNAs mapping to each virus in figure 2, separated by strand and 5'-base.

### S11 Repeatability of small RNA size distributions and base composition

The bar plots show the size distribution of small RNAs (17-29nt) for selected viruses (columns) separated across the 14 different sequencing libraries (rows). Bars plotted above the x-axis represent reads mapping to the positive strand and those below represent reads mapping to the negative strand. Bars are coloured according to the proportion of reads with each 5'-base (A-green, C-blue, G-yellow, U-red). The approximate number of reads for each virus in each pool is shown inset, and only viruses that had >100 small RNA reads in that pool are shown. Pairs of rows show technical replicates, and are labelled by metagenomic pool (E, I, K, S, T) or pool-mixture. For mixed pools (blue background) the reads in the lower rows were sequenced by Edinburgh Genomics following an oxidation step (to reduce miRNA representation), and the upper rows by BGI without an oxidation step. For unmixed pools (white background) the reads in the lower rows were sequenced using a ‘High Definition’ ligation protocol (both without oxidation). Note that the relative number of longer (23-27nt) reads seen in DCV, Nora Virus, Kilifi Virus and Thika Virus appears to be reduced in the presence of an oxidation step, as is the number of Kallithea miRNA reads. Raw count data for this figure are provided in S13.

### S12 Read count data for Figure 3

Raw counts of 21-23nt and 25-29 nt reads mapping to each de novo contig in Figure 3. The contigs are provided in supporting information S5.

### S13 Read count data for S11

Raw counts of 17-29nt small RNAs mapping to each virus in supporting information S11, separated by strand, 5'-base, and sequencing library.

### S14 Small RNA base composition by position

Part A ‘sequence logo’ plots show biased base composition along the length of virus-derived small RNAs, where the relative letter sizes indicates the base frequency among reads at that position, and the total height represents the information content (which quantifies bias away from equal frequencies). For each virus, only the most common read length is shown, and reads are combined across all sequencing libraries. Note that most viruses show a small bias toward A and U (plotted as T) at the 5' base, while Twyford Virus is biased toward U at both 5' and 3' positions. Kallithea DNA virus 22nt reads are dominated by the miRNA ATAGTTGTAGTGGCATTAATTG. Part B ‘sequence logo’ plots show biased base composition amongst viRNA-genome mismatches in 22nt and 23nt viRNAs from Twyford Virus (KP714075), indicating that many of the 3'U residues are non-templated. Raw count data for this figure are provided in S15.

### S15 Read count data for S14

Raw counts of bases for 21, 22, 23 and >24nt small RNAs mapping to each virus contig virus in supporting information S14.

### S16 Small RNA size and base composition for siRNA candidate viruses

The bar plots show the size distribution of small RNAs (17-29nt) for those unnamed siRNA-candidate loci that have >80 small RNA reads. Inset is the total number of small RNA reads summed across all libraries. Bars plotted above the x-axis represent reads mapping to the positive strand, and bars below the x-axis represent reads mapping to the negative strand. Bars are coloured according to the proportion of reads with each 5'-base (A-green, C-blue, G-yellow, U-red). All peak at 21nt and show the expected 5'-base composition (a slight bias against G), except for siRNA Candidate 14 (KP757950) which shows the 22-23nt 5' U-rich peak seen in Twyford Virus (KP714075). Count data are provided in S17.

### S17 Read count data for S16

Raw counts of 17-29nt small RNAs mapping to each siRNA candidate virus contig virus in supporting information S16, separated by strand, 5'-base, and sequencing library.

### S18 Relative small RNA production

To quantify differences in the rate at which small RNAs are generated from different viruses, we calculated the relative viRNA production for each of 16 different viruses in the ST and EIK metagenomic sequencing pools. The bars show the ratio between the relative number of 21-23nt small RNAs mapping to each virus (normalised by number of reads of the abundant *Drosophila* miRNA miR-34-5p), and the number of virus RNA-seq reads (relative to non-viral RNAseq reads, excluding rRNA reads). Ratios were then normalised to the lowest rate (Kilifi Virus in sample EIK) to give relative rates, such that 10^4^-fold more viRNAs are derived from DMelSV than from Kilifi Virus. Where viruses were present in both the EIK and ST pools, the correlation between the two datasets is high (rank correlation coefficient >0.99), suggesting that this variation is repeatable. Normalised data are provided in S19.

### S19 Relative count data for S18

The relative number of 21-23nt small RNAs mapping to each virus (normalised by number of reads of the abundant *Drosophila* miRNA miR-34-5p), and the number of virus RNA-seq reads (relative to non-viral RNAseq reads, excluding rRNA reads). Ratios were then normalised to the lowest rate (Kilifi Virus in sample EIK)

### S20 Evidence for a micro-RNA in Kallithea Virus

Output from the software package mirDeep [75] showing the proposed pre-miRNA hairpin, read numbers, and predicted folding pattern and energy. Reads are summed across all small-RNA libraries.

### S21 Read count data for Figure 4

The table contains normalised counts (viRNAs per million reads) of virus-derived small RNAs from 62 publicly available small-RNA sequencing projects (downloaded from the European Nucleotide Archive on 9^th^ May 2015). Up to 2 million reads (forward read only, if paired) were mapped for each sequencing run, and averaged across runs within projects.

### S22 Potential binding sites for Kallithea Virus miRNA

Potential miRNA binding sites in *D. melanogaster* 3' UTRs predicted by miRanda, and Gene Ontology enrichment analyses for genes with a miRanda score >150.

### S23 Virus-derived reads in publicly available RNA datasets

The grid shows the proportion of reads from each of 3144 publicly available RNAseq and small RNA datasets from *Drosophila* and *Drosophila* cell culture (vertical axis) that map to viruses (horizontal axis). Only datasets that have at least one virus present at ≥100 viral reads per million total reads were included, and only viruses present in at least one dataset at ≥100 viral reads per million total reads were included. Note that different parts of the segmented viruses generally co-occur within datasets, which allowed us to provisionally associate siRNA-candidate sequences with BLAST-candidate sequences (e.g. Nodaviruses and Bloomfield Virus). The viruses from DAV to *Drosophila melanogaster* Birnavirus were reported previously, the others are newly described here. Full data are presented in S24, and exemplar cell culture datasets are presented in S26.

### S24 Read count data for Figure 4 and S26

The table contains counts of virus-derived small RNA and RNA-seq reads from 9656 publicly available sequencing ‘run’ datasets (downloaded from the European Nucleotide Archive on 9^th^ May 2015). Up to 2 million reads (forward read only, if paired) were mapped for each dataset. Datasets with no mappable reads were excluded.

### S25 Virus-like sequences identified by *de novo* assembly of public reads

Virus-like contigs assembled by *de novo* from publicly available *D. melanogaster* RNAseq datasets (which cannot be submitted to Genbank), including Brandeis Virus, Berkeley Virus, a Totivirus-like sequence and several reovirus-like sequences.

### S26 Viruses in widely-used *Drosophila* cell cultures

A single exemplar RNAseq dataset was selected for each of 26 widely-used *Drosophila* cell culture lines, and all forward reads were mapped to viruses. Twenty four of the datasets were drawn from ModEncode cell culture sequencing [81], and two that were not available (OSS and Kc167) were taken from datasets SRR070269 and SRR500470. The colour scale illustrates the fraction of reads that were viral in origin (virus reads per million total reads) and viruses are sorted in ascending order of viral read fraction (from ML-DmD32 with < 1% viral reads, to ML-DmBG1-c1 with ∼50% viral reads). Note that the virus population is likely to differ between different subcultures, and these values should only be considered illustrative. Raw counts data for these datasets are provided in S24.

### S27 Survey collection details, locations and virus prevalence

Collection data (sample sizes, city, country, latitude and longitude, date) and virus prevalence for each species at each location (maximum likelihood estimate with 2 log-likelihood confidence intervals, all rounded to 2 sig. fig.).

### S28 Virus prevalence by population

Charts show virus and *Wolbachia* prevalence in *D. melanogaster* and *D. simulans* for each of the 17 locations at which flies were collected. Values are maximum likelihood estimates with 2 log-likelihood intervals, and are plotted on a log scale. Where no bar or confidence interval is provided, no flies of that species were sampled. Location details and raw data are provided in **S27**.

### S29 Rates of multiple infection and co-infection

Panels (A) and (B) show the percentage of *D. melanogaster* and *D. simulans* carrying at least one of the surveyed viruses. Bars are are maximum-likelihood estimates with 2 log-likelihood intervals. Coloured bars represent carriers of DAV, DCV, Nora, DMelSV, Twyford, La Jolla, Kallithea, Dansoman, Torrey Pines, Craigie’s Hill, Thika, Newfield, Bloomfield or Motts Mill viruses (but excluding the high-prevalence siRNA-candidate viruses), grey bars show the large increase in overall prevalence if the siRNA-candidate viruses (Galbut Virus and Chaq Virus) are included. Missing bars indicate that the host species was absent from the collection location (for collection details see S27). Panels (C) and (E) show the proportion of virus-free and multiply-infected *D. melanogaster* when Galbut Virus and Chaq Virus are excluded (C) or included (E), while (D) and (F) show the equivalent plots for *D. simulans*. Panels (C-F) are calculated using single-fly assays only (not bulks), and are averages across populations.

### S30 Correlation between *Wolbachia* and virus prevalence

Each plot shows the correlation between the named virus and *Wolbachia* in prevalence, across the sampled populations. Inset are rank correlation coefficients, and a simple linear regression (virus ∼ *Wolbachia*) for illustration. The analysis does not account for any spatial autocorrelation in prevalence, and does not correct for multiple testing. Points are coloured by location (see S28 and S29 for colours) and plotted with 2 log-likelihood intervals. Nominal ‘significance’ at *p*<0.05 is shown using asterisks. Data are provided in S27.

### S31 Evolutionary properties of *Drosophila* viruses

Each panel shows parameter estimates of viral evolution for the eleven RNA viruses for which sufficient sequence data were available. Panels are (A) mutation rate, (B) date for the most recent common ancestor, (C) the range of movement between geographic regions (defined as continents and laboratory), (D) the rate of switching between *D. melanogaster* and *D. simulans*, and (E) the relative rates of synonymous (*dS*) and nonsynonymous (*dN*) substitution (*dN-dS*>0 implies positive selection). For panels A-D points are the median of the posterior sample and 95% credibility intervals, panel E shows the median and range across codons. For all viruses except DAV and Nora Virus, a strong prior was placed on mutation rate (Panel A, grey box), and mutation rates and MRCA dates were inferred separately for each alignment block. The underlying BEAST xml files (including alignments and model specifications) are provided, along with the resulting mcc tree files and summaries of posterior distributions, in supporting information S32 and S33. The underlying FUBAR batch files and per-site parameter estimates are provided in supporting information S34.

### S32 Alignments and BEAST files for Figure 6 and S31

Alignments and BEAST models as BEAST xml for Figure 6 and S31

### S33 Maximum clade credibility trees for Figure 6 and S31

Posterior summaries of BEAST output for Figure 6 and S31 (posteriors inferred jointly across two independent runs).

### S34 FUBAR analysis and output for Figure 6 and S31

FUBAR (HyPhy) command batch files and output. Alignments were those provided in S32, and trees were those provided in S33.

### S35 *dN/dS* in DAV and Nora Virus

Point estimates of *dN/dS* are shown for each codon in open reading frames of DAV and Nora Virus (ratios of the posterior estimates, not the posterior estimates of the ratios). Bar colours illustrate posterior support that the site is constrained (blue for strong support that *dN*< *dS*) or positively selected (red for strong support that *dN>dS*). Positions coloured grey have little support for either positive selection or constraint. The dashed horizontal line indicates neutrality (*dN=dS*), so that bars higher than this are candidate sites for positive selection. The solid horizontal line shows the mean of the *dN/dS* estimates for that open reading frame. *dN/dS* estimates greater than 3 have been truncated to 3 for clarity. FUBAR batch files and parameter estimates are provided in supporting information S34.

